# Rapid adipose tissue expansion triggers proliferation of memory adipose tissue T cells and accumulation of novel T helper 2 (Th2) regulatory cells

**DOI:** 10.1101/2025.10.22.683711

**Authors:** Ramiah D. Jacks, Peizi Wu, Jennifer L. Delproposto, Carey N. Lumeng

**Affiliations:** Department of Pediatrics, University of Michigan Medical School, Ann Arbor, MI

**Keywords:** Adipose tissue T cells, Obesity, Th2 cells, HFD, Transcriptomics, Adipose Tissue Inflammation

## Abstract

Obesity generates a pro-inflammatory state contributing to immune dysregulation and metabolic disease. In response to overnutrition, adipose tissue immune cells undergo dynamic changes featuring a shift from a type 2 anti-inflammatory phenotype in lean adipose tissue to a pro-inflammatory phenotype in obese adipose tissue, contributing to obesity associated adipose tissue inflammation featuring an accumulation of macrophages and T cells. Adipose tissue T cells (ATTs) from mice and humans with obesity have an exhausted phenotype, suggesting chronic activation. However, the mechanism(s) that regulate the balance between anti-inflammatory and pro-inflammatory ATT phenotypes with obesity are not understood. We sought to investigate how and if ATT cells respond to early obesity. We characterized the temporal and phenotypic changes in ATTs in response to initial adipose tissue remodeling, using short-term high fat diet (stHFD) as a model. We observed the accumulation of ATTs in the visceral adipose tissue (VAT) due to the induction of proliferation in memory, and not naïve, ATTs with 7 days of HFD. Cellular Indexing of Transcriptomes and Epitopes by sequencing (CITE-seq) demonstrated a decrease in Tregs and γδ T cells with an increase in a novel Th2 cell population with immune regulatory capacities featuring expression of the immunoregulatory cytokine *Tgfb1,* and an activated gene expression profile featuring an increase in *Nr4a1* and *Itk*, in VAT with 14 days of HFD. This study represents the first description of a Th2 regulatory cell population in adipose tissue, and its increase with short-term HFD suggests underexplored mechanisms of ATT subpopulation activation and regulation.

**GRAPHICAL ABSTRACT:** 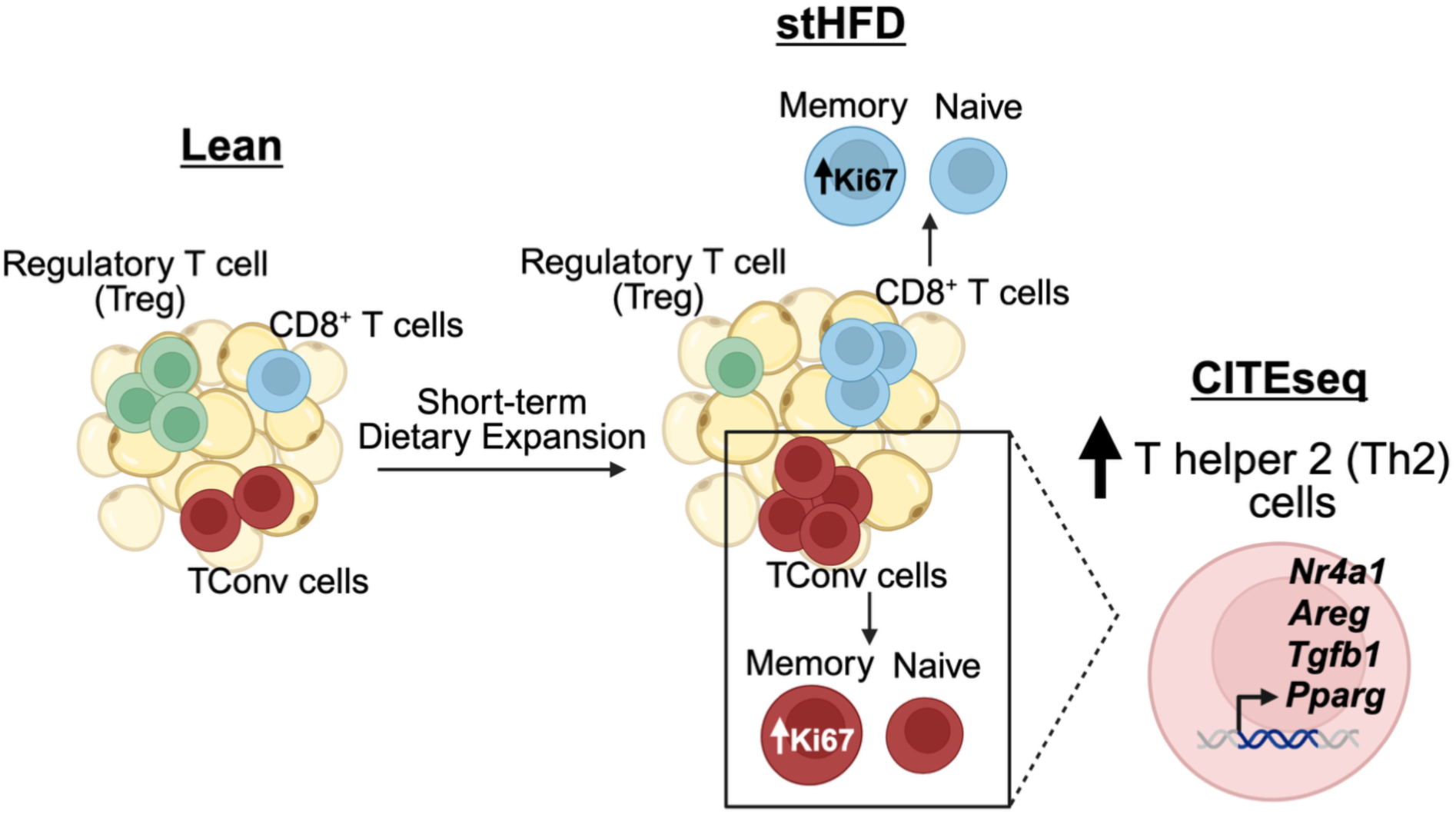

## 1. INTRODUCTION

Over 890 million adults live with obesity worldwide, corresponding to 1 in 8 people. Obesity is a risk factor for the majority of the leading causes of death including heart disease, stroke, chronic obstructive pulmonary disease, lung cancer, diabetes, kidney disease, fatty liver disease, and chronic liver disease[1]. A feature of obesity is chronic adipose tissue inflammation that is associated with metabolic disease and adipose tissue dysfunction and is regulated by a network of resident adipose tissue leukocytes[2]. In response to overnutrition, adipose tissue immune cells undergo dynamic changes[3]. The first and second most prevalent immune cell populations in adipose tissue, adipose tissue macrophages (ATMs) and adipose tissue T cells (ATTs) shift from an anti-inflammatory phenotype in lean adipose tissue to a pro-inflammatory phenotype in obese adipose tissue, contributing to obesity associated adipose tissue inflammation[4].

Resident ATTs in lean adipose tissue are comprised of CD4^+^ T helper 2 (Th2) cells and regulatory T cells (Tregs) whose main functions are to maintain an anti-inflammatory environment and tissue homeostasis[4, 5]. Flow cytometry, single cell RNA-sequencing studies and Cellular Indexing of Transcriptomes and Epitopes by sequencing (CITE-seq), demonstrate that these anti-inflammatory ATTs decrease with chronic obesity with a significant increase in pro-inflammatory T helper 1 (Th1) cells and CD8^+^ T cells[6, 7]. Diet induced obesity (DIO) mouse models deficient in Th1 cells or CD8^+^ T cells exhibited reduced adipose tissue inflammation and improved glucose sensitivity compared to wild type mice, providing evidence for pro-inflammatory ATT involvement in promoting obesity associated inflammation[8, 9].

The mechanisms that activate and reshape ATT and systemic T cell dysfunction are not completely understood. MHC-II signals from ATMs and adipocytes have been implicated in ATT cell activation[10–12] Clonal TCR rearrangements have been described in ATT cells suggesting antigen dependent processes shape the ATT cell pool[13]. ATT cell exhaustion, a state of effector T cell functional impairment with reduced capacity to secrete cytokines and concurrent increased expression of inhibitory receptors, has been described in obese adipose tissue[14],[6, 15]. T cell exhaustion is induced by chronic antigen stimulation, suggesting ATT activation/stimulation within the adipose tissue microenvironment with obesity. In people with obesity, peripheral T cell impairment may contribute to susceptibility to infection and impaired vaccine responses, as observed during the COIVD-19 pandemic[16–19].

Since the factor(s) and mechanism(s) that regulate the balance between anti-inflammatory and pro-inflammation ATT phenotypes with obesity are not understood, the goal of our study was to investigate how and if ATT cells respond to early obesity associated with rapid expansion and remodeling of adipose tissue. Elucidation of such mechanism(s) can potentially identify targets to suppress detrimental pro-inflammatory ATT accumulation and rescue anti-inflammatory ATTs to maintain metabolic homeostasis. To address this, we characterized the temporal and phenotypic changes in ATTs in response to initial adipose tissue remodeling, using short-term high fat diet (stHFD) as a model. Our previous data demonstrate that stHFD, as defined as 7 days (7d) and 14 days (14d) of feeding in mice, results in rapid expansion of adipose tissue mass, adipocyte hypertrophy, and a concurrent increase in resident ATMs due to proliferation[20]. Using this model, we investigated if primary ATT responses are also induced in this dynamically and rapidly changing adipose tissue microenvironment. Similar to ATMs, we observed the accumulation of ATTs in the visceral adipose tissue (VAT) due to the induction of proliferation amplified in memory, and not naïve, ATTs with 7d of HFD. CITE-seq demonstrated a decrease in Tregs and γδ T cells with an increase in a novel Th2 cell population with immune regulatory capacities in VAT with 14d of HFD. This study represents the first description of a Th2 regulatory cell population in adipose tissue, and its increase with short-term HFD suggests underexplored mechanisms of ATT subpopulation activation and regulation.

## 2. MATERIALS AND METHODS

### 2.1. Mice

Male and female C57BL/6J (000664) mice were purchased from Jackson Labs at 17 weeks of age. Male and female OT-II mice (004194) were purchased from Jackson Labs and bred in house. At 18 weeks of age, mice were fed ad libitum a HFD (60% fat, Research Diets, D12492) or normal chow diet (13.4% fat, LabDiet PicoLab, 5L0D) for 7 and 14 days. All studies were approved by the Institutional Animal Care and Usage Committee of the University of Michigan.

### 2.2. Stromal vascular fraction and splenocyte isolation

Mice were euthanized by isoflurane overdose followed by cervical dislocation. eWAT, iWAT, and spleen were collected and weighed. Adipose tissue depots were minced and digested in 10 mL of 1 mg/mL type II collagenase (Sigma-Aldrich) for 30 minutes at 37°C. After digestion, adipose tissue was filtered through 100 μm filters, RBC lysed and filtered through 70 um filters as previously described[15]. Spleens were crushed on a 70 uM filter and RBC lysed.

### 2.3. Flow Cytometry

The stromal vascular fraction (SVF) and splenocytes were incubated in Fc Block (eBioscience) at 1:100 for 5 minutes at 4°C. Next, cells were extracellularly stained and stained for viability for 30 minutes at 4°C. Anti-mouse extracellular antibodies included: CD45, CD11b, CD3, CD4, CD8, CD44, CD62L, and Live Dead Violet. Cells were washed with FACS buffer and fixed and permeabilized using a Foxp3 transcription factor staining buffer kit (eBioscience) for 45 minutes at room temperature. Cells were washed with permeabilization buffer and intracellularly stained with anti-mouse Foxp3 and Ki67 for 45 minutes at room temperature. More information on the antibodies used for flow cytometry can be found in Supplemental Table 1. Cells were analyzed on a Cytek Aurora Spectral Flow Cytometer and data was visualized and analyzed with Flow Jo Software (Tree Star Inc).

### 2.4. CITEseq and Immune Profiling

SVF cells from individual mice (5, lean normal diet fed and 5, 14 day HFD fed) were incubated in Fc block (eBioscience) for 5 minutes at 4°C. Next cells from individual mice were stained with unique hashtag antibodies (1:200) (Biolegend). Biological replicates were pooled and CD45^+^ immune cells were enriched using CD45 microbeads (Miltenyi Biotec), LS columns (Miltenyi Biotec), the QuadroMACS Separator (Miltenyi Biotec).

Collected CD45^+^ SVF cells were then stained for CITE-seq using TotalSeq-C mouse universal cocktail according to the manufactures instructions and stained for ATT sorting using Live Dead Violet and the following anti-mouse antibodies: CD45, CD3, CD11b for 30 minutes at 4°C. Cells were sorted on a BigFoot Cell Sorter (Thermo Fisher Scientific) with the sorting gate set to collect Live CD45^+^ CD3^+^ CD11b^-^ ATTs for submission for sequencing. Anti-mouse antibodies used for sort were a different clone that those same targets in the TotalSeq-C mouse universal cocktail prevent competition for binding. A complete list of the hashtag antibodies, antibodies used for sorting and the 113 antibodies contained within the TotalSeq-C mouse universal cocktail can be found in Supplemental Table 2 and Supplemental Table 3.

Samples were submitted to the University of Michigan’s Advance Genomics Core for library construction using the single cell 5’ workflow with the Immune Profiling kit for the 10X Chromium platform (10X Genomics) and Illumina sequencing (10B NovaSeq). We targeted 10,000 cells per diet group and a sequencing depth of 10,000 reads per cell.

### 2.5. Data processing

The R package Seurat V5 was used for quality control, normalization, integration, clustering, and subclustering analysis[21]. For quality control, cells were filtered to retain those with ≥200 and ≤1,500 detected gene features and ≤5% mitochondrial transcript content. Furthermore, cells being identified as doublets based on hashtag demultiplexing (HTOs) were excluded from downstream analysis. The gene counts (RNA) were normalized using SCTransform[22] and further regressed out mitochondrial read percentage. Dimensionality reduction of the RNA modality was performed via reciprocal principal component analysis (rPCA). To correct for batch effects across 10 biological samples, we applied Harmony on the rPCA embeddings. ADT (antibody-derived tag) assays were normalized using centered log-ratio (CLR) transformation, followed by principal component analysis (PCA) for dimensionality reduction. A joint representation of the RNA and proteomic modalities was constructed using weighted nearest neighbor (WNN) analysis, leveraging Harmony-corrected RNA embeddings and CLR-transformed ADT PCA embeddings.

### 2.6. Annotating clusters

Clustering was performed using the smart local moving (SLM) algorithm implemented in Seurat, with a low resolution (0.3) to identify broad cell populations. Differentially expressed markers were identified using a non-parametric Wilcoxon rank-sum test, with P-values adjusted for multiple testing using the Bonferroni correction. Clusters annotation was guided by cell type markers and surface markers, and further supported using SingleR V2.4.1[23] with the Immgen[24] database. Clusters identified as CD4+ were further subset into individual samples, and re-integrated using the method mentioned above. Subclusters were annotated based on the expression of marker genes and surface proteins.

### 2.7. Downstream sequencing analysis

Comparison across clusters and dietary groups were performed using the SCTransfom assay with a non-parametric Wilcoxon rank sum test. Gene set enrichment analysis (GSEA) was carried out using package fgsea[25] V1.28.0, utilizing gene sets from Hallmark collection, Gene Ontology (GO) Biological Processes, Kyoto Encyclopedia of Genes and Genomes (KEGG), and Reactome pathway databases. Package scRepertoire[26] V2.0.4 was used for T cell receptor (TCR) analysis and visualization. For pseudotime and trajectory inference, Monocle3[27] V1.3.4 was used.

## 3. RESULTS

### 3.1. ATTs accumulate with ATT subtype and depot specific differences in response to rapid adipose tissue expansion

To determine if stHFD effects ATTs dynamics, we fed male C57BL/6 mice HFD (comprised of 60% fat) *ad libitum* for 7 days (7d) or 14 days (14d) or fed a normal chow diet (ND). Total body weight was significantly increase with 7d of HFD feeding and remained increased with 14d of HFD feeding (Figure 1A). Both epidydimal (eWAT) and inguinal (iWAT) white adipose tissue depots increased rapidly with HFD, with eWAT more than doubling by 14d (Figure 1B). Total number of live stromal vascular fraction (SVF) cells significantly increase in the eWAT, but not iWAT, with 7d and 14d HFD (Figure 1D). However, upon normalization to fat pad weight, there is no significant difference in the cells per gram in the eWAT, suggesting that the increase in cell numbers is proportional to the increase in fat pad weight (Figure 1C). To determine if there are any changes in ATTs with stHFD, we assessed the frequency and cell number of CD45^+^ leukocytes and CD3^+^ ATTs by flow cytometry (Figure 1D). While the frequency of CD45^+^ leukocytes and CD3^+^ ATTs remained comparable at all timepoints, the quantify of CD45^+^ cells per fat pad were significantly increase with HFD in the eWAT, but not the iWAT (Figure 1E). Further, CD3^+^ ATT cell numbers were significantly increased with 14d of HFD only in the eWAT (Figure 1E). Taken together, these data suggest that stHFD induces an increase in CD3^+^ ATT cells in the eWAT, but not iWAT.

**Figure 1.**
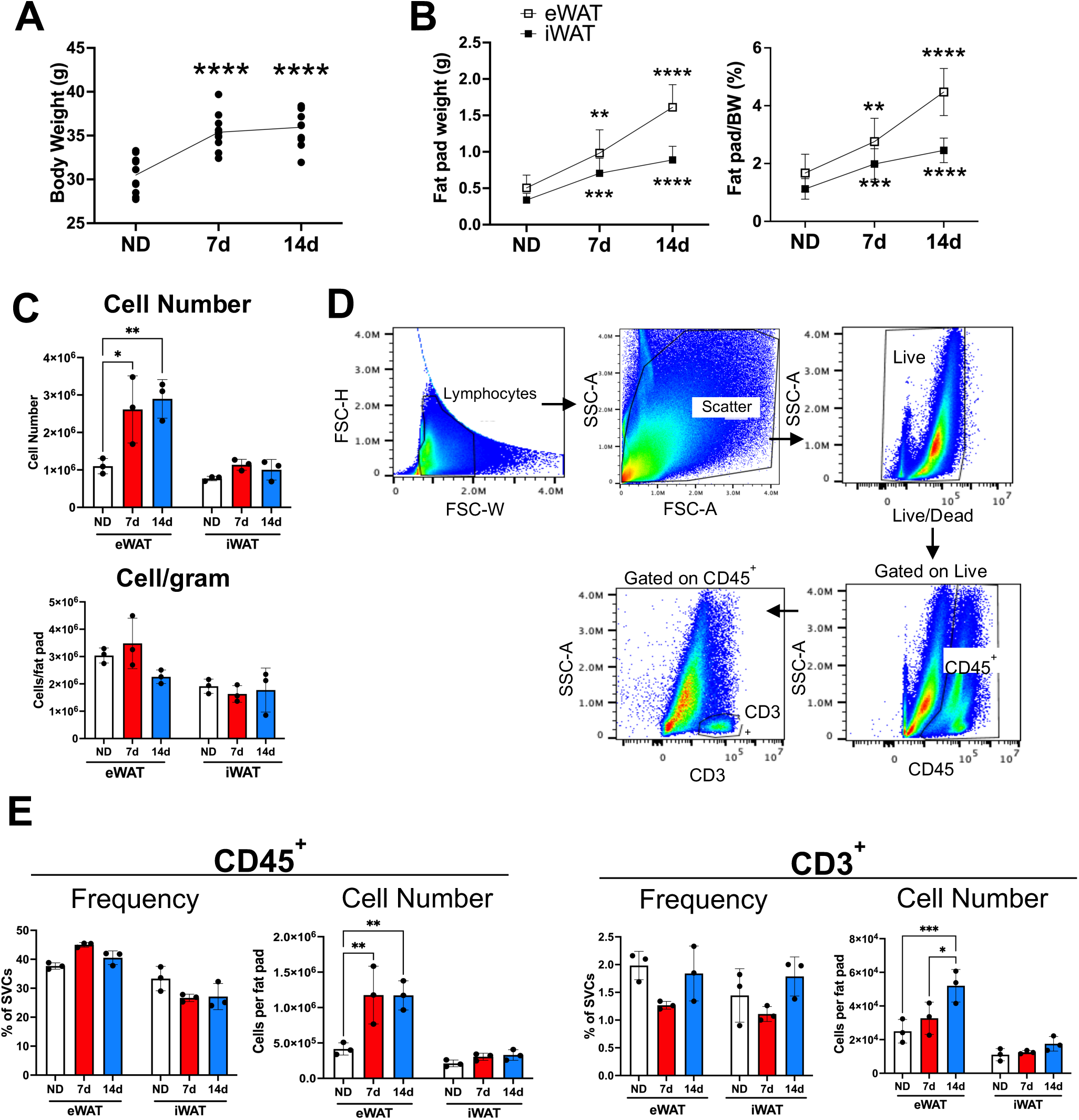
Short-term high-fat diet promotes rapid white adipose tissue expansion and increase adipose tissue T cell numbers in male mice. Total body weights (A) and fat pad weights and fat pad weights normalized to body weight (BW) (B) of mice fed a normal diet (ND) or 60% high-fat diet for 7 or 14 days. (C) Representative flow plots showing gating strategy for assessing CD3^+^ adipose tissue T cells (ATTs). (D) Quantity of stromal vascular fraction cells (SVCs) in eWAT and iWAT reported as total cell numbers and normalized to fat pad weight. (E) Frequency and quantity of CD45^+^ and CD3^+^ ATTs in mice fed a normal diet (ND) or 60% high-fat diet for 7 or 14 days. *n* = 3 mice/group. 3 independent experiments. Analyzed by one-way ANOVA with Tukey’s multiple comparisons where ** *P* < 0.01, *** *P* < 0.001, and **** *P* < 0.0001.

There are several types of T cells that reside within lean adipose tissue, including CD4^+^ regulatory T cells (Tregs), CD4^+^ conventional T cells (Tconv) which includes CD4^+^, Foxp3^-^ T helper (Th) cells), and CD8^+^ T cells. To determine which ATT cell types are responding to stHFD, we assessed the frequency and quantity of each type by flow cytometry. We observed a significant decrease in Treg frequency within the CD4^+^ T cells with 7d and 14d of HFD in eWAT without significant increases in quantity per fat pad between ND and 14d in the eWAT (Figure 2B). Tregs were not altered in iWAT (Figure 2B). Conversely, there is a significant increase in the frequency of conventional Tconv in the eWAT and cell number of these cells in the eWAT and iWAT with 14d of HFD (Figure 2B). Finally, while there is an increase in the frequency of CD8^+^ T cells with 7d, this increase is not maintained at 14d in the eWAT (Figure 2B). We observed a significant increase in CD8^+^ T cells quantity in the eWAT and iWAT with 14 days of HFD (Figure 2B). These data demonstrate that the increased accumulation of ATTs in the eWAT at 14d (Figure 1E) can be attributed to expansion of the pool of Tconv and CD8^+^ T cells.

**Figure 2.**
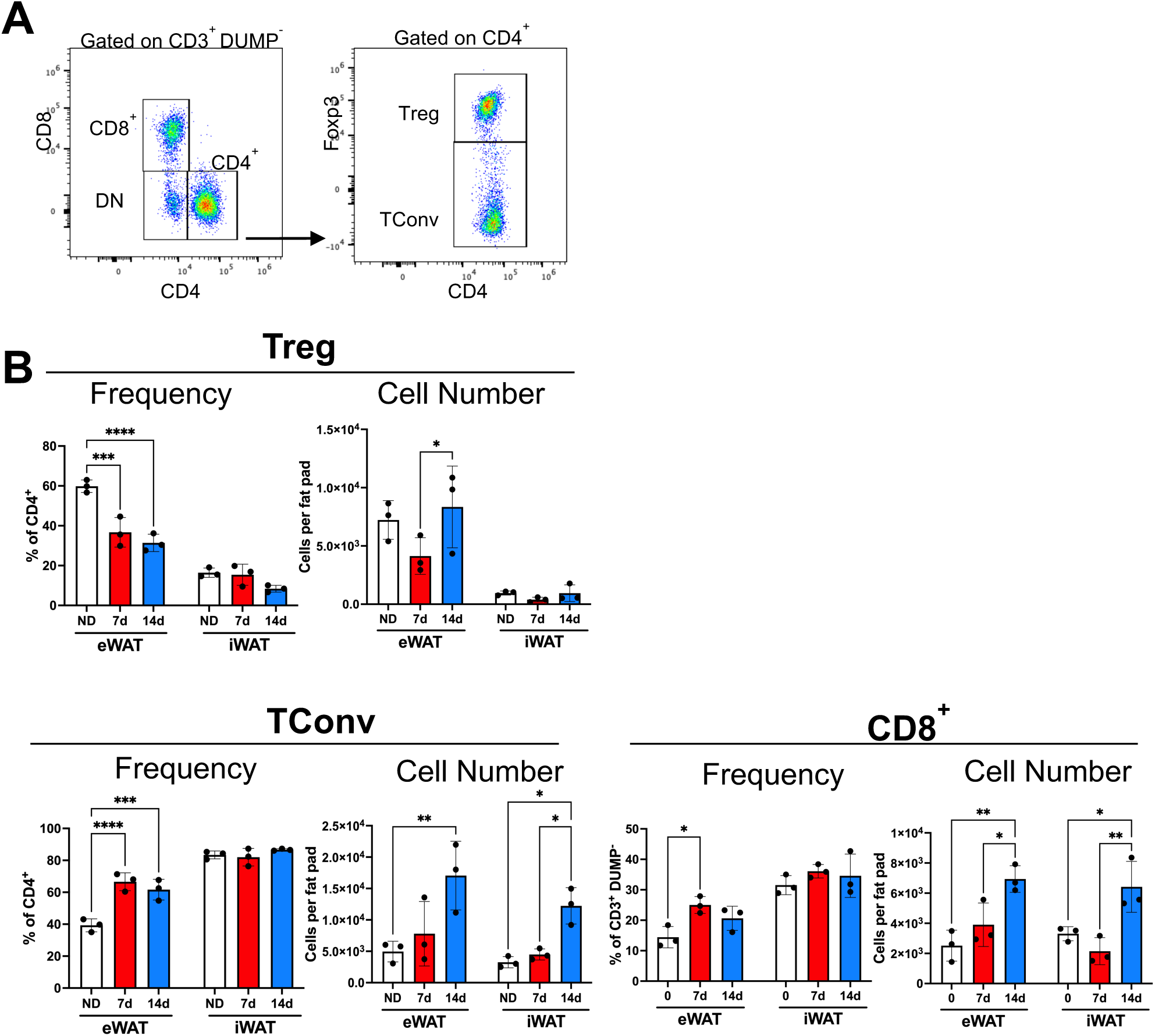
Adipose Tissue T cell subsets increase in number with short-term high-fat diet. (A) Representative gating strategy to assess CD4^+^ regulatory T cells (Tregs), CD4^+^ conventional T cells (Conv T), CD8+ and double negative (DN) ATTs. (B) Frequency and quantity of ATT subsets in mice fed a normal diet (ND) or 60% high-fat diet for 7 or 14 days. *n* = 3 mice/group. 3 independent experiments. Analyzed by one-way ANOVA with Tukey’s multiple comparisons where ** *P* < 0.01, and *** *P* < 0.001.

Further, these data indicate that stHFD decreases the frequency of Tregs in the eWAT, but not in the iWAT, highlighting depot specific differences in ATT regulation and that ATTs cell populations are remodeled in quickly upon initiation of HFD induced obesity.

### 3.2. stHFD induces memory ATT proliferation

We next sought to understand the mechanisms responsible for the increase in Tconv and CD8^+^ T cells. We hypothesized that ATTs proliferate in response to stHFD to contribute to the increase in cell numbers. To test this, we assessed proliferation of ATTs at 7d and 14d of HFD. We observed a significant increase in proliferating (Ki67^+^) T cells with 7d of HFD in Tregs and Tconv in the eWAT and iWAT, with no change in proliferation in CD8^+^ T cells (Figure 3B). We further hypothesized that this observed proliferation in response to stHFD was specific to ATT, as we previously published that T cell exhaustion in response to HFD was only observed in ATT, and not in T cells from the spleen[15]. To test this, we examined proliferation in splenic T cells spleen and observed no difference in proliferation with 7d or 14d of HFD in Treg, Tconv or CD8^+^ T cells (Figure 3B). The induction of proliferating ATT cells was most prominent at 7d. At 14d of HFD ATT proliferation returned to levels comparable to ND suggesting that the induction of ATT proliferation in CD4^+^ Tregs and Tconv occurs soon after initiation of HFD in eWAT and iWAT.

**Figure 3.**
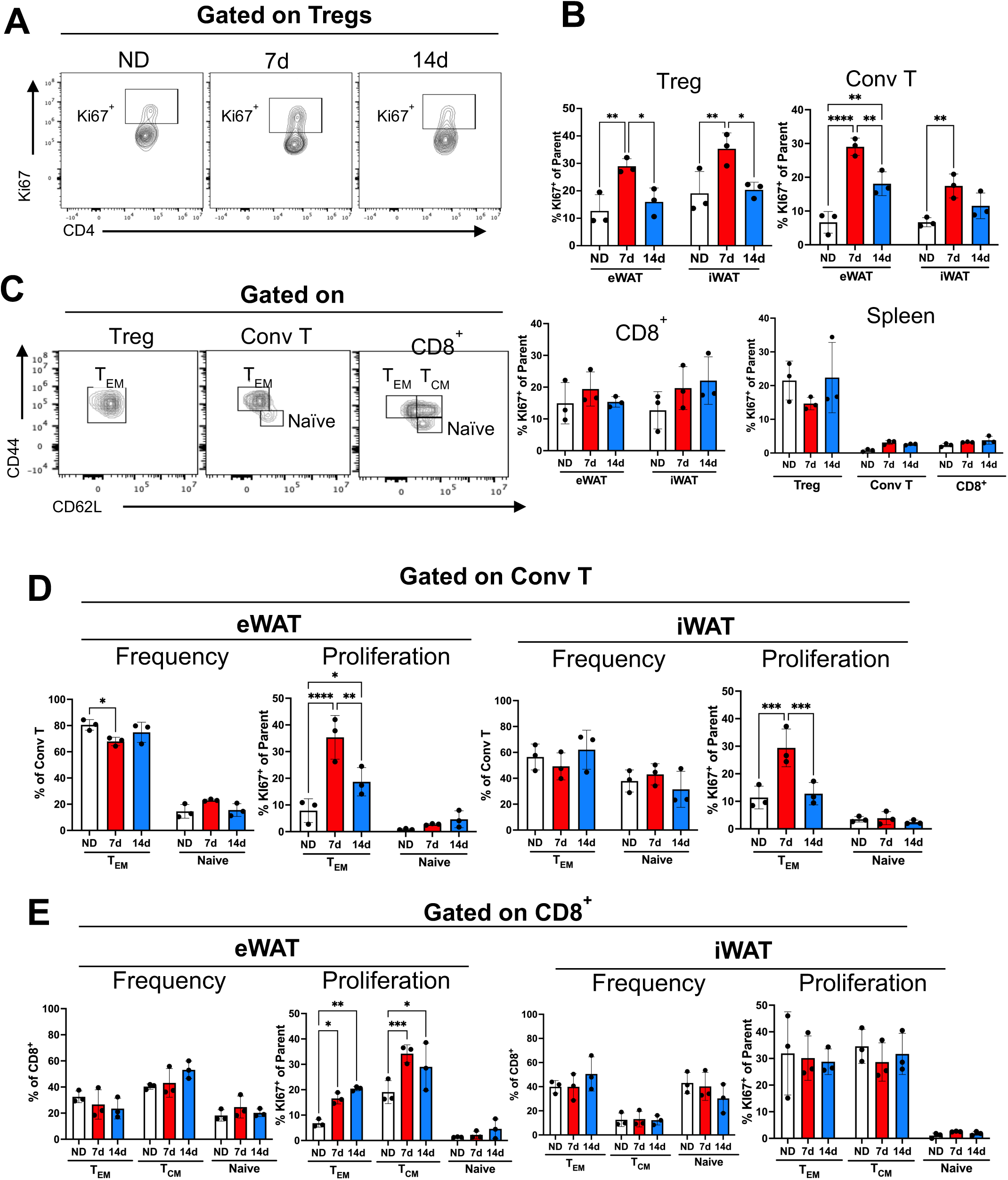
Adipose Tissue T cells proliferate with 7 days of high-fat diet. (A) Representative gating strategy to assess proliferation by ki67 expression in mice fed a normal diet (ND) or 60% high-fat diet for 7 or 14 days. (B) Ki67 expression in ATT subsets in eWAT, iWAT and spleen. (C) Representative gating strategy to assess CD44^+^ CD62L^-^ effector memory (T_EM_), CD44^-^ CD62L^+^ naïve, and CD44^+^ CD62L^+^ central memory (T_CM_) ATTs within ATT subsets. Frequency and proliferation of memory and naïve subtypes within conventional T ATTs (conv T) (D) and CD8^+^ ATTs (E) in eWAT and iWAT. *n* = 3 mice/group. 3 independent experiments. Analyzed by one-way ANOVA with Tukey’s multiple comparisons where ** *P* < 0.01, *** *P* < 0.001, and **** *P* < 0.0001.

We further sought to determine if proliferation we observed in ATTs with stHFD was occurring in memory or naïve ATTs. Memory T cells are antigen-experienced and retain the memory of previous antigen activation and have the ability to rapidly respond to the same antigen to which they were generated[28]. Conversely, naïve T cells are antigen inexperienced, and upon antigen activation they can differentiate into a variety of T cell subsets[29]. Consistent with prior reports, we observed that CD4^+^ and CD8^+^ ATTs are largely comprised of effector memory T cells (T_EM_; CD44^+^ CD62L^-^) with ∼60% (iWAT) to 80% (eWAT) for CD4^+^ Tconv and ∼60% in eWAT and iWAT for T_EM_ and central memory (T_CM_; CD44^+^ CD62L^+^) combined in CD8^+^s (Figure 3D and 3E)[11]. In Tconv, an increase in proliferating ATTs is induced with 7d of HFD specifically in the T_EM_ ATTs population in both the eWAT and iWAT. No proliferation was observed in naïve T cells (Figure 3D).

Moreover, in CD8^+^ ATTs, we observe two subtypes of memory T cells, effector memory (T_EM_) and central memory (T_CM_) in adipose tissue (Figure 3E). We observed an increase in proliferating T_EM_ and T_CM_ with 7d and 14d of HFD, but not naïve T cells, in the eWAT (Figure 3E). We observed no induction of proliferation with stHFD in CD8^+^ memory subtypes in iWAT (Figure 3E). These data suggest that stHFD induces the expansion of memory ATTs by inducing proliferation.

### 3.3. ATT accumulation with stHFD is not observed in female mice

It is well appreciated that there are sex differences in the development of obesity and insulin resistance in mice and humans[30–32]. Indeed, less obesity-associated adipose tissue inflammation is observed in female mice and women compared to male mice and men[33]. Further, female C57BL/6 mice exhibit reduced fat pad weight gain and reduced insulin resistance with diet-included obesity models compared to male mice[33]. We sought to determine if ATT responses to stHFD were different in female mice compared to male. Female C57BL/6 mice were fed HFD for 7d and 14d and observed a significant increase in total body weight with 14 days and an increase in gonadal WAT (gWAT) and iWAT with 7 and 14 days of HFD (Figure 4A and B). The ∼2 fold increase in gWAT mass at 14d in females was less than the ∼3 fold increase in eWAT mass at the same timepoint observed in males. There was no significant increase in total cell number, CD45^+^ immune cells, nor CD3^+^ ATTs in females (Figure 4C and D). Despite the lack of increase in ATT cell numbers, we observed a significant increase in proliferating Tregs with 7d of HFD in female gWAT, but not in the iWAT (Figure 4E) which contrasted with what we observe in males (Figure 3B). Further, we observed a significant increase in proliferation in Tconv T_EM_ with 7d of HFD in both the gWAT and iWAT (Figure 4F) that was similar to that observed in males. In female mice, we did not observe a significant induction of proliferation with HFD in T_EM_ nor T_CM_ in CD8^+^ ATTs in the gWAT and iWAT compared to ND (Figure 4E). Overall, while we observed an induction of CD4^+^ Tregs and Tconv T_EM_ proliferation in response to stHFD in females, in contrast to males, this did not translate into an increase in total ATT cell number.

**Figure 4.**
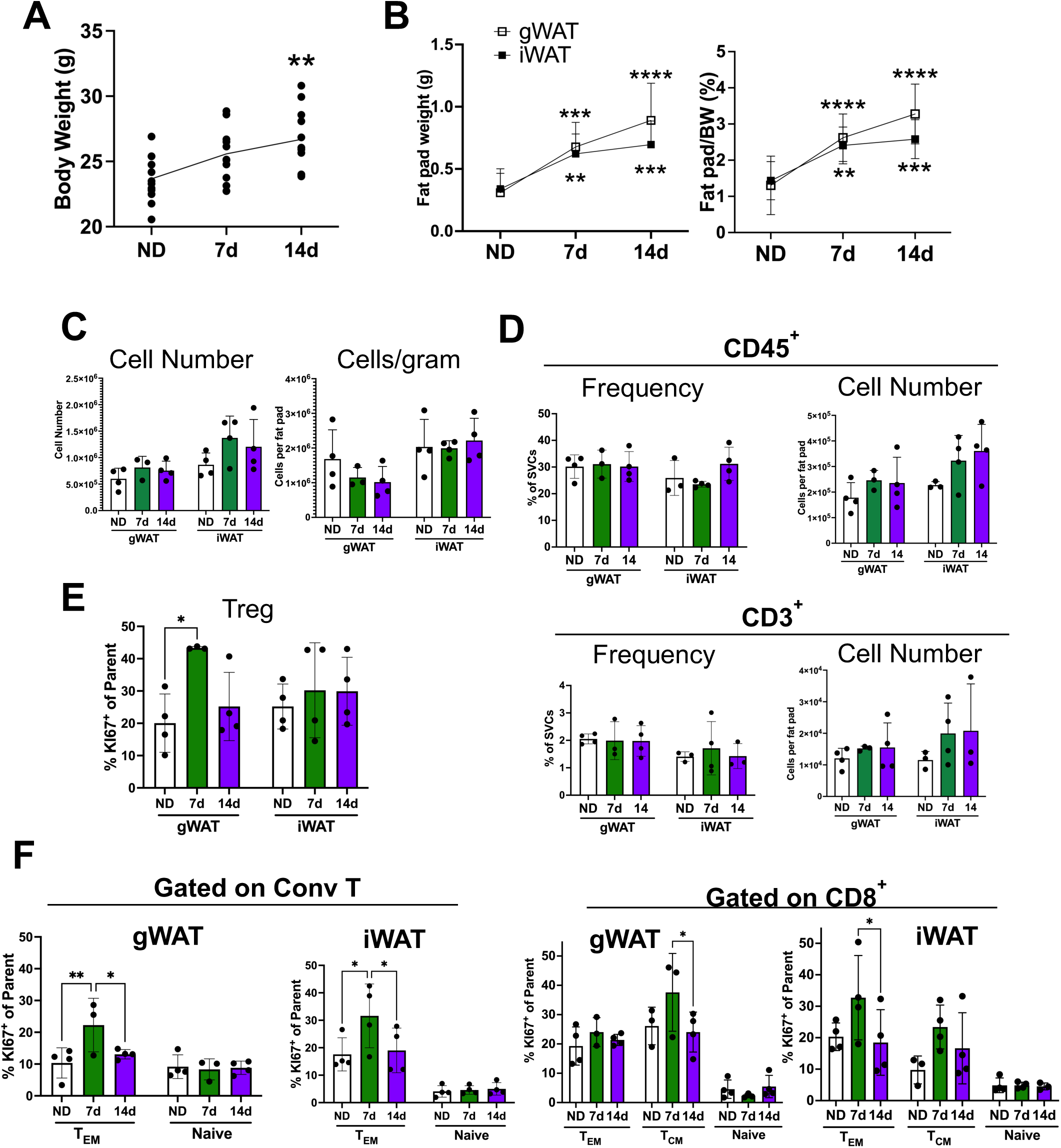
Short-term high-fat diet promotes rapid white adipose tissue expansion with no increase adipose tissue T cell numbers in female mice. Total body weights (A) and fat pad weights and fat pad weights normalized to body weight (BW) (B) of female mice fed a normal diet (ND) or 60% high-fat diet for 7 or 14 days. (C) Quantity of stromal vascular fraction cells (SVCs) in gWAT and iWAT reported as total cell numbers and normalized to fat pad weight. (D) Frequency and quantity of CD45^+^ and CD3^+^ ATTs in mice fed a normal diet (ND) or 60% high-fat diet for 7 or 14 days. *n* = 3 mice/group. (E) Ki67 expression in Tregs in gWAT and iWAT. (F) Ki67 expression of memory and naïve subtypes within conventional T ATTs (conv T) and CD8^+^ ATTs in gWAT and iWAT. 3 independent experiments. Analyzed by one-way ANOVA with Tukey’s multiple comparisons where ** *P* < 0.01, *** *P* < 0.001, and **** *P* < 0.0001.

### 3.4. CITE-seq shows that stHFD suppresses Tregs and γδ and activates Th2 ATT cells

We next sought to address two major remaining questions that arose from these data. First, we were interested in understanding the nature of the stimulation that drives the ATT proliferation we observed with stHFD. Second, we wanted to understand the phenotype and function of the ATTs that respond to stHFD stimuli to understand how/if these ATT participate in the orchestration of the adipose tissue inflammatory state we observe with obesity. To address these questions, we employed CITE-seq along with T Cell Receptor (TCR) sequencing. CITE-seq allows to the acquisition of surface protein data and gene expression data simultaneously, which assists in the delineation of T cell subtypes and activation markers[34]. We enriched for CD45^+^ immune cells from the eWAT of male mice fed a ND or a HFD for 14d, FACS sorted CD3^+^ CD11b^-^ ATTs, and performed CITE-Seq with immune cell profiling (TotalSeq-C, Supplemental Figure 1).

UMAP plots revealed 10 ATT subsets (Supplemental Figure 1 B, C, and D). We manually annotated these ATT subsets by integrating antibody and gene expression data in the following manner: Natural Killer (NK) T cells (NK1.1), CD8^+^ memory (CD8, CD44), CD8^+^ naïve (CD8, CD62L), T helper 1 (Th1) CD4^+^ (CD4, CD44, *Ifng*), CD4^+^ proliferating (CD4, CD44, *Ki67*), CD4^+^ naïve (CD4, CD62L), T helper 2 CD4^+^ (CD4, CD44, *gata3*), CD4^+^ *Ifng*-memory (CD4, CD44), Tregs (CD4, CD44, *Foxp3*), and γδ T cells (TCR γδ, *Cd163l1*) (Supplemental Figure 1 C and D, Supplemental Table 4 and 5).

Comparing the proportions of ATTs between ND with 14d HFD mice, the most prominent changes observed were a decrease in Tregs and γδ T cells and an increase in CD4^+^ T helper 1 (Th1) cells and CD4^+^ T helper 2 (Th2) cells (Figure 5A and B). To understand cell-type specific gene expression changes, gene set enrichment analysis (GSEA) of Tregs revealed a downregulation of pathways involved in T cell activation in Tregs from HFD fed mice compared to ND (Figure 5C, Supplemental Table 6 and 7). The most significantly downregulated genes in the T cell activation pathway in Tregs include *Ltb4r1* (leukotriene B4 receptor 1) whose expression in activated T cells promotes migration to inflammatory sites[35], *Dusp1* (map kinase protein 1, MKP-1) a positive regulator of T cell proliferation and activation[36], *Klrg1* (killer cell lectin-like receptor G1) whose expression is associated with activated and memory Tregs[37], *Cd48* whose surface expression is associated with T cell activation[38] and *Tigit*, a co-inhibitory receptor associated with the suppressive function of Tregs[39] (Figure 5C). These data suggest that stimuli from short-term HFD feeding reduces Treg frequency and their activation/suppressive phenotype.

**Figure 5.**
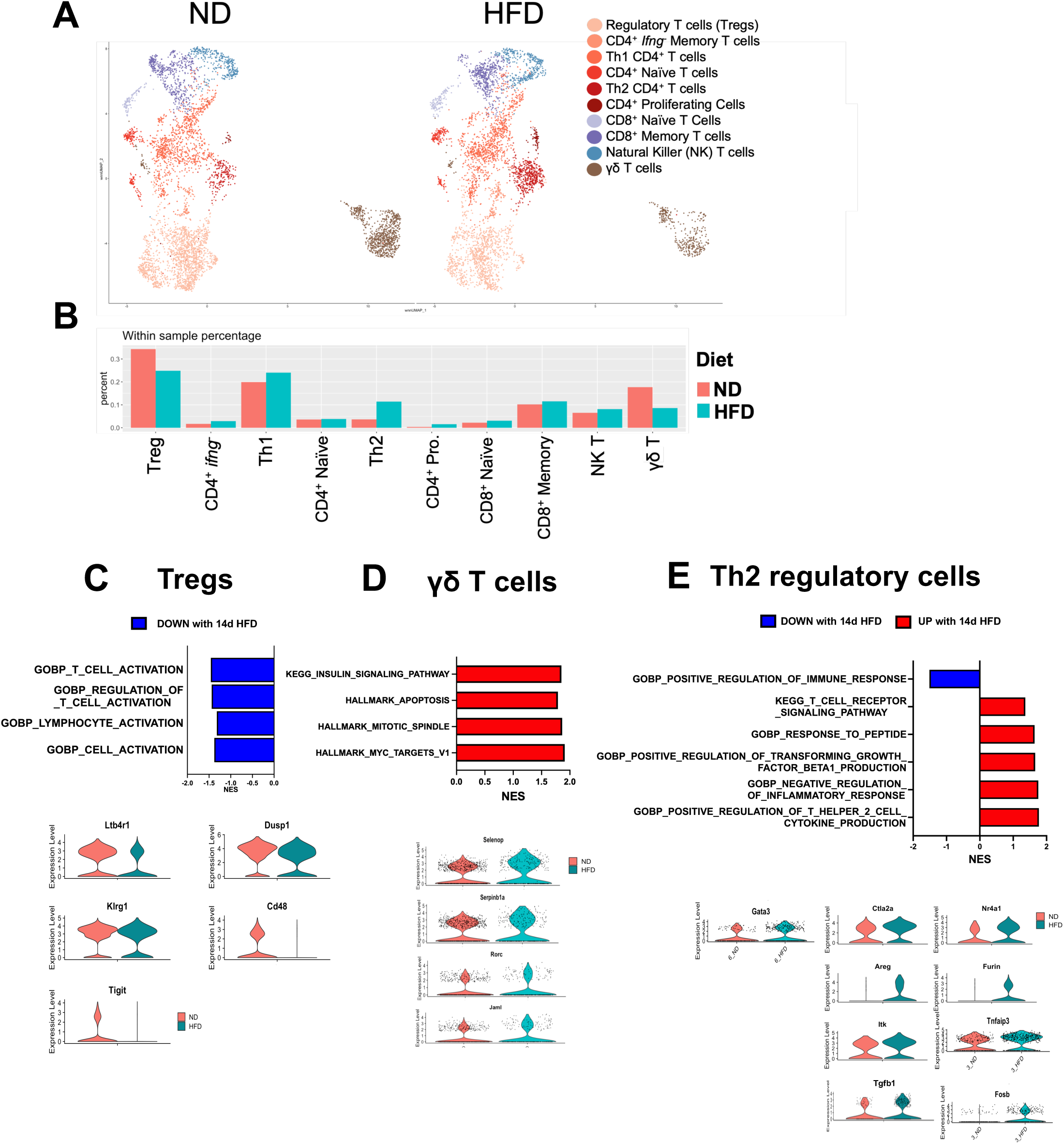
Adipose Tissue T cell changes with 14 days of high-fat diet. (A) UMAP between ND and HFD samples of ATTs from the eWAT of mice fed a normal diet (ND) or a high-fat diet (HFD). (B) Quantification of ATT subpopulation changes between ND and HFD. (C-E) Pathway analysis and top differentially expressed (DE) genes in Tregs, γδ T cells, and Th2 regulatory cells.

While γδ T cells have been shown to increase in cell number in chronic diet-induced obesity mouse models[40], we observed a dramatic decrease in γδ T cells with 14 days of HFD (Figure 5B). We observed an increase in pathways related to proliferation (MYC targets V1 and V2, mitotic spindle), an increase in the insulin signaling pathways, and apoptosis pathways in γδ T cells with 14 days of HFD (Figure 5D, Supplemental Table 8). These data suggest that signals within the adipose tissue microenvironment might promote apoptosis of γδ T cells during short-term HFD.

The most prominent observation was an increase in Th2 cells with 14d HFD. Genes involved in the negative regulation of inflammatory response pathways were upregulated and positive regulation of immune response was downregulated with stHFD.(Figure 5E, Supplemental Table 9). The negative regulation of inflammatory response pathway includes *Furin*, a protease that cleaved inactive transforming growth factor beta (TGF-β) into its active form[41]. Further, positive regulation of TGF-β production was upregulated with stHFD, and in increase in *TGFβ* gene expression with stHFD, corresponding with the increase in *Furin.* Additionally, T cell activation and proliferation pathways including the T cell receptor signaling pathway, response to peptide, Th2 cell cytokine production pathways were also upregulated with stHFD in Th2 cells (Figure 5C). We observed a significant increase in genes associated with T cell activation/TCR stimulation with stHFD, including *Nr4a1*[42], *Itk*[43, 44], *Fosb*[45], and *TNFaip3*[46] as well as cytotoxic T lymphocyte-associated protein alpha 2 (*ctla2a)*, whose expression is observed in activated T cells but its function in negative immune regulation from T cells isn’t fully elucidated [47]. These data suggests that Th2 cells become activated and may possess regulatory functions that have not been previously elucidated in adipose tissue Th2 cells.

To validate our observations of Th2 ATT induction with stHFD, we performed t-SNE analysis on our flow data to increase the resolution for ATT cell subtypes. Flow cytometry confirmed the suppression of Tregs with stHFD. We also observed an increased in CD8 mem T cells. Consistent with the CITE-seq data, we observed an increase in a CD4^+^ CD44^+^ CD62L^-^ Ki67^+^ Foxp3^-^ with markers consistent with Th2 ATTs (Supplemental Figure 2).

### 3.5. Th2 regulatory ATT shares a gene expression profile with canonical Th2s and Tregs

To better delineate Th2 and Tregs, we performed a subset analysis on CD4^+^ ATTs which recapitulated the observation of a decrease in Tregs (Figure 2B) and demonstrated an increase in Th2 cells with HFD (Figure 6A and B, Supplemental Tables 10-11). Since our data indicate that Th2s can have regulatory functions, we compared their gene expression profile with Tregs. (Figure 6C) In terms of genes known to be associated/shared with both Tregs and adipose tissue Th2s, both Th2 and Tregs express *Gata3, Areg (*amphiregulin), and *Il1rl1* (ST2/IL-33 receptor). In addition, both cell types express *Tgfb1,* with a substantial portion of the Th2 population expressing *Tgfb1*, and *Pparg,* which has been previously implicated as an important regulator of ATT Treg function and accumulation. (Figure 6C)[48–50]. ATT Th2 cells also express the canonical type 2 cytokines *Il4* and *Il5*[51]. Finally, ATT Tregs express *Foxp3,* the signature transcription factor associated with Tregs. Taken together, these data suggest that stHFD induces the accumulation of eWAT Th2 ATTs featuring gene expression of canonical Th2 associated genes as well as the immunoregulatory cytokine *Tgfb1*.

**Figure 6.**
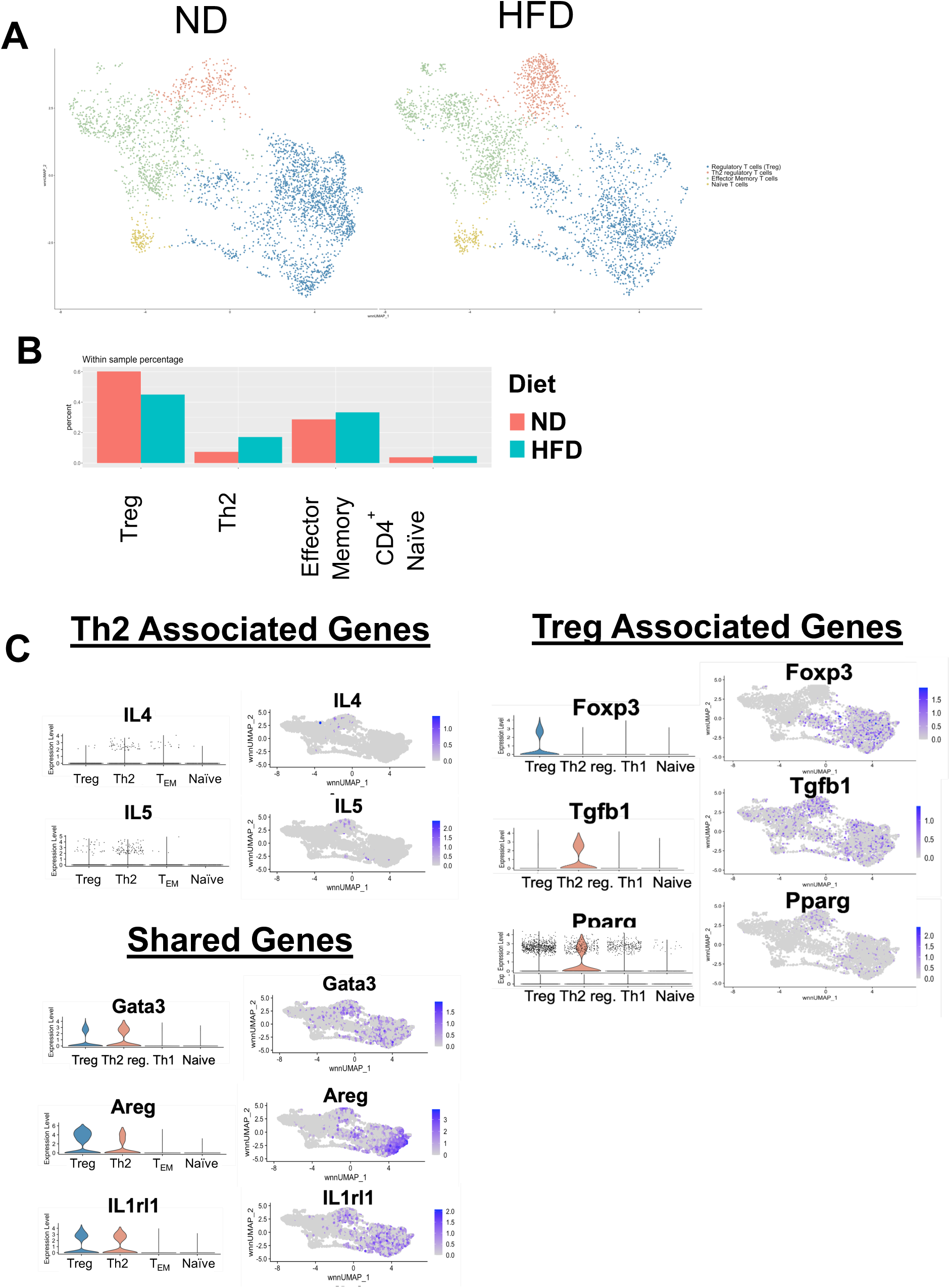
Th2 regulatory cluster shares gene expression prolife with Tregs. (A) Uniform Manifold Approximation and Projection (UMAP) generated by gating on CD4^+^ ATTs only need to know how to describe this better. (B) CD4^+^ ATTs UMAP between ND and HFD samples of ATTs from the eWAT of mice fed a normal diet (ND) or a high-fat diet (HFD). (C) Quantification of ATT subpopulation changes between ND and HFD. (D) Gene expression profiles of genes associated with Th2 cells and Tregs.

### 3.6. Th2 regulatory ATTs with unique TCR sequences increase with stHFD

TCR sequencing was used to identify clonal populations in the CITE-seq dataset. We identified TCR clones in all CD4^+^ ATT subsets in both dietary conditions that ranged in size from 1 cell to 69 cells in the ND and 38 cells in HFD (Figure 7A). The largest number of unique TCR sequences as a proportion of the subsets were observed in the CD4^+^ T_EM_ and Treg clusters (Supplemental Figure 3B). With stHFD, there is a slight decrease in number of unique TCR sequences in both CD4^+^ T_EM_ and Treg ATTs (Supplemental Figure 3B). In contrast, stHFD led to an increase in unique TCR sequences in the Th2 cell subset (Figure 7B). No specific clonal sequences were observed in the ND samples that were expanded by stHFD in any CD4^+^ ATT cluster and there was no overlap between clones we identified in the ND samples and clones in the HFD samples (Figure 7C). Taken together, that stHFD induces expansion of unique TCRs in Th2 ATTs, but not Tregs or T_EM._

**Figure 7.**
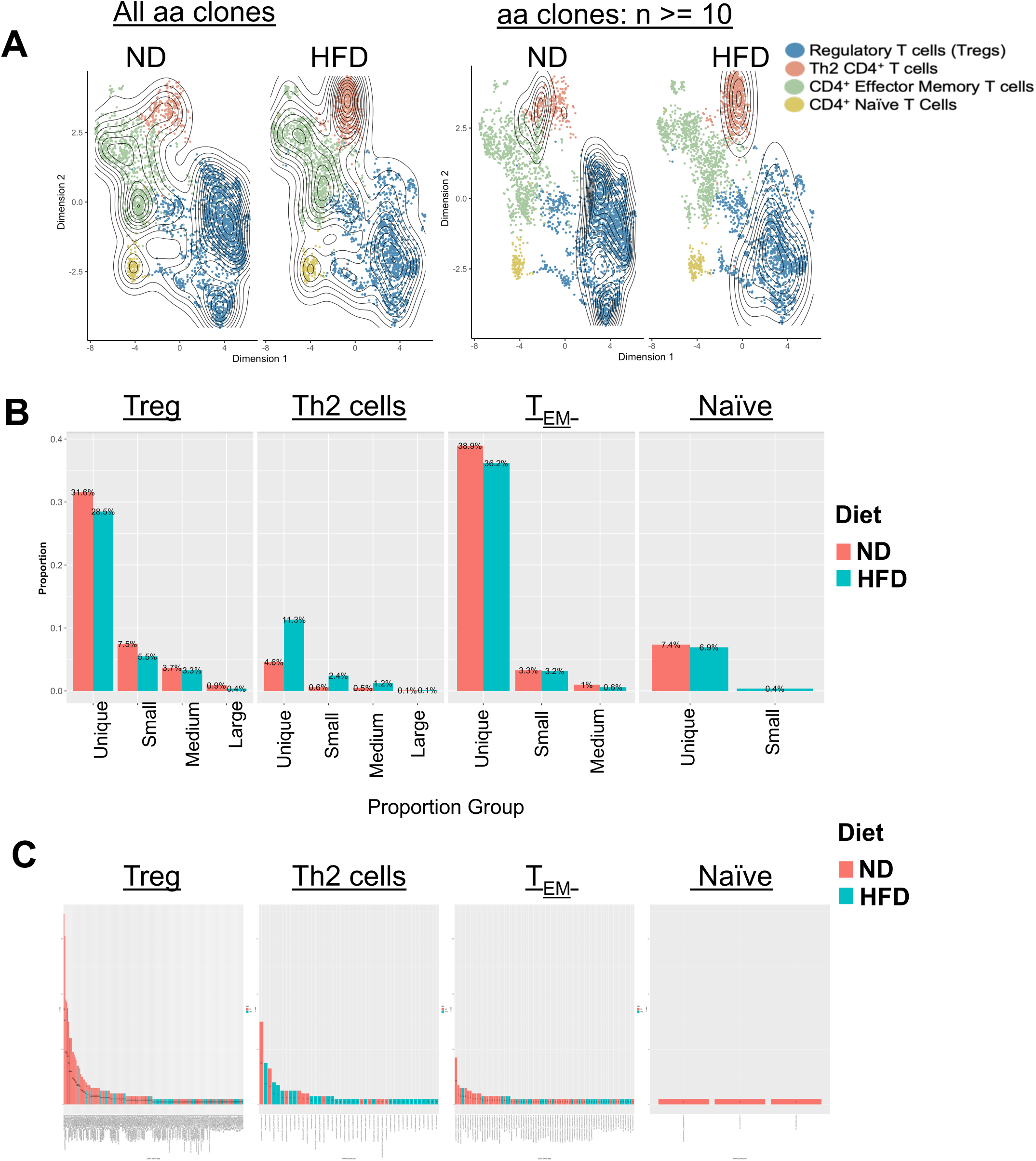
Th2 regulatory cluster is clonally diverse with 14 days of HFD. (A) TCR clone mapping of all clones (left) and TCR clones greater than 10 (right) in UMAP of CD4^+^ ATTs. (B) TCR clone sequence and frequency in Tregs, Th1 cells and Th2 regulatory ATTs with ND and 14 days of HFD. Unique x = 1, small (0.001 < p < = 0.002), medium (0.002 <p<= 0.01), large (0.01 < p < = 0.1). (C) Number of cells per TCR sequence in ND and HFD CD4^+^ ATT subpopulations.

The shared genes between Th2 cells and Tregs (*Areg, Il1rl1*, *Pparg*) suggest that they may share a common lineage or are ex-Tregs, which are Tregs that lose the expression of *Foxp3*[52]. To assess this, we evaluated the distribution of clones with >5 cells identified between the CD4^+^ ATT subsets. Clonal populations identified in the Th2 population were also present in the T_EM_ subsets and were generally not shared with Tregs. For example, in the largest clone of 47 ATTs from a ND mouse, 30 of those cells are Th2 and 17 are T_EM._ In contrast, Treg clones were largely unique to the Treg pool (Supplemental 3A top). Integrating individual mouse data allowed for a more directly lineage linkage, as all 47 cells within the largest clone are from one mouse (ND 2) (Supplemental 3A bottom). The clonal analysis suggests that Th2 ATTs are derived from Tconv and not Tregs. Evidence for this also comes from single-cell trajectory analysis (Supplemental 3B), that supports a differentiation trajectory for Th2 cells that branches from T_EM_ and not Tregs. Overall, our data suggests that Th2 ATTs induced by stHFD are indeed terminally differentiated Th2s and are not derived from Tregs.

### 3.7. Tregs and CD4^+^ T_EM_ expressing TCRVβ5^+^ proliferate in response to stHFD

To evaluate the role of antigen-specific signals in ATT expansion with stHFD, we employed an antigen specific anti-OVA T cell receptor (OT-II) mouse model with an OVA-specific TCR comprised of TCRVα2^+^ and TCRVβ5^+^. Some CD4^+^ T cells in OT-II mice do not have the OVA-specific TCRVα2^+^ TCRVβ5^+^ TCR enabling us to distinguish antigen specific and non-specific signals (Figure 8B)[53]. OT-II mice were fed HFD for 7d and we observed significantly greater proliferation in eWAT OT-II^+^ Tregs and CD4^+^ T_EM,_ and iWAT Tregs compared to OT-II^-^ Tregs and CD4^+^ T_EM_ within OT-II mice (Figure 8C). Since we didn’t inject the mice with OVA, we reasoned that maybe one singular TCR chain might be important for responding to HFD challenge, instead of the combination of TCRVα2^+^ TCRVβ5^+^ that confers OVA specificity. To test this, we examined the TCR composition of cells within C57BL/6 mice and asked if there is a difference in the proliferation of T cells based on their expression of TCRVβ5^+^ and/or TCRVα2^+^ with 7d of HFD (Figure 8D). We observed that proliferation was induced with 7d of HFD in eWAT and iWAT Tregs and CD4^+^ T_EM_ ATTs with TCRVβ5^+^ single positive and TCRVα2^+^ TCRVβ5^+^ double positive TCRs, and not ATTs with TCRVα2^+^ and double negative TCRs (Figure 8E). These data suggest that the expression of TCRVβ5 might be important in responding to HFD challenge to drive ATT proliferation.

**Figure 8.**
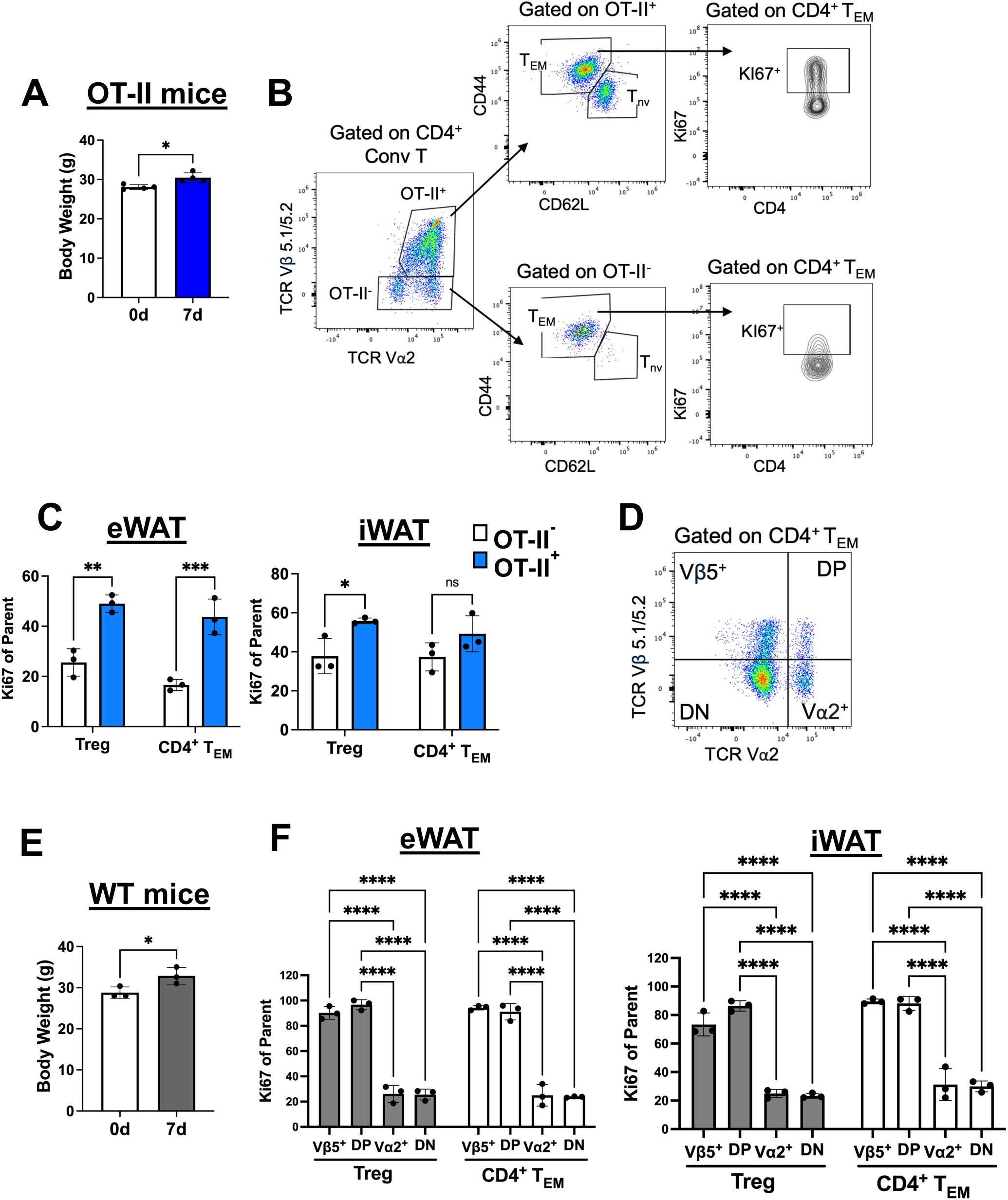
Adipose Tissue T cell proliferation is observed in OVA specific and non-specific memory T cells. Male OT-II mice were placed on a 60% high-fat diet for 7 days. (A) Body Weight in OT-II mice (B) Representative gating strategy to assess TCRVα2^+^ TCRVβ5^+^ OT-II^+^ and TCRVα2^-^ TCRVβ5^-^ OT-II^-^ CD4^+^ ATTs. (C) Ki67 expression in Tregs and effector CD4^+^ conventional ATTs (CD4^+^ T_EM_) in the eWAT and iWAT. (D) Representative gating strategy to assess TCR composition as TCRVβ5^+^ TCRVα2^-^, TCRVβ5^+^ TCRVα2^+^ (double position, DP), TCRVβ5^-^ TCRVα2^+^, and TCRVβ5^-^ TCRVα2^-^ (double negative, DN). (E) Body wight in wild type (WT) mice. (F) Ki67 expression in Tregs and CD4^+^ T_EM_ in eWAT and iWAT stratified by TCR composition. Analyzed by one-way ANOVA with Tukey’s multiple comparisons where ** *P* < 0.01, *** *P* < 0.001, and **** *P* < 0.0001.

## 4. DISCUSSION

Numerous ATT subpopulation changes occur during short-term HFD further demonstrating that dynamic adipose tissue changes are associated with immune stimulation. Previous studies investigating early ATT kinetics with HFD show an increase number of CD8^+^ ATTs with 14 days of HFD, and an increase number of macrophages with 6 weeks of HFD in the eWAT, suggesting a role for ATT in macrophage recruitment in visceral adipose tissue[54]. Flow cytometry data presented here echo the increased number of CD8^+^ ATTs and additionally demonstrate an increased number of CD4^+^ conventional ATTs with 14 days of HFD. Obese adipose tissue features a dramatic elevation CD8^+^ ATTs and conventional CD4^+^ ATTs, namely, Th1s. Our data highlight that some of the changes we observe in adipose tissue with obesity are already beginning to develop with just 14 days of HFD. We see that ATTs respond to early alterations in the adipose tissue with HFD, however, the precise mechanism remains under intense investigation. A more recent study proposed that adipose tissue stem cells (ASC) secrete C-C motif chemokine ligand 5 (CCL5) with 4 weeks of HFD that enhances peripheral T cell recruitment infiltration into adipose tissue[55]. Conversely, work demonstrated that resident memory ATTs proliferate with only 7 days of HFD and we hypothesize that the increase ATT frequency we observe by flow cytometry is due to the proliferation of resident memory ATTs, and not infiltration of peripheral T cells. This is probable, due to the timing difference between the recorded infiltration of peripheral T cells at 4 weeks and our study examining up to 2 weeks. Our data and others show that memory ATTs comprise the majority of T cells in adipose tissue, with upwards of 80% of ATT being of the memory phenotype. Memory ATTs have been shown to participate in protective immune responses against pathogens, providing evidence that adipose tissue is an immunological organ[3, 56]. Therefore, we theorize that the mechanism(s) that induce memory ATT proliferation early in response to HFD might be different than the mechanisms required for peripheral T cell recruitment into adipose tissue. Further, the differentiation and establishment of memory ATTs requires further examination to determine their roles in orchestrating inflammation in adipose tissue early in the development of obesity.

We previously demonstrated depot specific ATT function dysfunction whereby ATT exhaustion was observed the VAT of mice and humans[15]. Work presented here show ATT accumulation in the VAT (eWAT) and not in the subcutaneous WAT (iWAT). These data highlight depot-specific differences in ATT regulation with HFD. Further, impaired insulin signaling and chronic low-grade inflammation is associated with VAT expansion in obesity[57]. Our data demonstrate that the drivers of ATT accumulation reside uniquely within VAT and may not be present, or are suppressed in SAT with adipose tissue expansion.

One striking feature of ATTs in obesity adipose tissue is the decreased frequency of Tregs in mine and humans[10, 58]. Not only are Tregs decreased with obesity as early as 8 weeks of HFD, their abundance does not recover with weight loss[6, 59]. Our flow cytometry and CITEseq data both reveal a decrease in Treg frequency with 14 days of HFD, much earlier that these previously published reports. Of note, however, we did not observe a decrease in Treg cell number within the timepoints studied (Figure 2B), suggesting that the decrease frequency is reflecting the comparison of Tregs to Tconv in that Tconv frequency significantly increases with short-term HFD (Figure 2C). Further, since our data show a significant increase in total CD3^+^ ATT numbers with 14d HFD, the absences of a change in Treg numbers suggests no expansion of resident Tregs compared to lean conditions. Several proposed mechanisms exist to explain the decrease in Treg numbers with obesity, including interferon-gamma (IFN-γ) included apoptosis, loss of cells that produce the Treg survival cytokine, IL-33, and decreased expression of the IL-33 receptor (*Il1rl1*)(ST2) in Tregs. Our CITEseq data in Tregs show a significant decrease in T cell activation pathways with 14d HFD. We did not observe an increase in apoptosis pathways or a statistically significant difference in expression of the IL-33 receptor (*Il1rl1*)(ST2) with 14d HFD, suggesting that induction of apoptosis or loss of the ability to sense the IL-33 survival signal may not be the mechanisms responsible for our observed decrease in Treg frequency. Further, *pparg* has been reported as a master regulator of the visceral adipose tissue Treg phenotype, including regulation of ST2 expression[50]. We observed no difference in *pparg* expression in Tregs with 14d HFD. Because our CITEseq only examined CD3^+^ ATTs, we do not know if there are early changes in the abundance of IL-33 secreting cells, such as Innate Lymphoid Cells type 2 (ILC2s), which are decreased with obesity. Based on our data, we speculate that Treg activation is altered or dysregulated with short-term HFD and other published methods of Treg reduction develop later with obesity. The precise T cell activation signaling pathways that could be suppressed in Tregs with short-term HFD remain to be elucidated.

Reported sexual dimorphism in obese mice feature VAT Tregs accumulation due to estrogen signaling, reduced adipose tissue inflammation, reduced fat pad weight gain, and reduced insulin resistance in female mice compared male mice[60, 61]. While we observed an increase in fat pad weights in female mice, they did not expand to the level as male mice. We also did not observe an accumulation of ATTs with HFD in female mice, as we saw in male mice. Interestingly we did observe some proliferation of Tregs and memory CD4^+^ and CD8^+^ ATTs with 7d HFD. We speculate that the observed proliferation does not translate into ATT accumulation this is due to documented differences in anti-inflammatory cell populations in female which function to limit immune cell expansion, which we are observing even with early adipose tissue expansion.

We observed a decrease in the abundance of γδ T cells with 14 days of HFD in the eWAT by CITEseq (Figure 5B). Previous studies have shown that γδ ATT cells accumulate in with 5 to 10 weeks of HFD [40]. Further, mice that are deficient in γδ T cells exhibited reduced adipose tissue inflammation and decreased systemic insulin resistance compared to γδ T cell sufficient mice after 10 to 24 weeks of HFD, suggesting the γδ T cells promote inflammation and insulin resistance [40]. Our study used a 60% HFD while other studies examining γδ T cell with obesity used 45% Western HF. Further, a ketogenic diet in mice was demonstrated to activate metabolically protective γδ T cell in VAT which restrains inflammation [62]. Taken together, these data suggest that diet may assist in driving specific γδ T cell outcomes. Future studies are required to determine if the decreased abundance of γδ T cells we observed at 14 days of 60% HFD is reversed at later timepoints.

Literature states that Th2s decrease in adipose tissue with obesity and their frequency in adipose tissue is inversely correlated with systemic inflammation and insulin resistance in humans[63]. While Th2s are proposed to aid in the maintenance of an anti-inflammatory environment through the secretion of the type 2 cytokines IL-4 and IL5, the specific roles and functions of Th2s in adipose tissue are understudied[4, 64]. Our CITEseq data demonstrated that Th2 increase in abundance with stHFD, indicating these cells respond to early change within adipose tissue. To date, our work provides the first evidence of a Th2 population in adipose tissue with an immunoregulatory phenotype. Strikingly, our CITEseq reveled that eWAT Th2s express *tgfb1* in lean and 14d HFD VAT, with a significant increase in expression with stHFD (Figure 5E). Further, TGF-β has been believed to be inhibitory for Th1 and Th2 differentiation and function while being essential for the differentiation of T helper 17 cells (Th17) and Tregs [65, 66]. TGF-β is a potent immunoregulatory cytokine that can inhibit the proliferation and function of immune cells and is a hallmark characteristic of Treg function. While *tgfb1* expression in Th2 in adipose tissue has not been recorded, a previous study reported that peripheral Th2 cells can be induce to secrete of active TGF-β in an helminth infection model, which serve an immunoregulatory function and was sufficient to control graft vs host disease (GVHD)[67]. The combinatory evidence of increased abundance of Th2s, increased pathways and genes associated with T cell activation and negative regulation of inflammatory response strongly suggest an immunoregulatory role for adipose tissue Th2s with stHFD. Further TGF-β is also implicated in regulating adipocyte biology, namely, differentiation and proliferation [68]. It remains undetermined if *tgfb1* expression in Th2s affects adipocytes.

Interestingly, these Th2 regulatory cells increased in abundance and appeared to be selectively activated with Tregs decreased with stHFD. Our data does not support the hypothesis that the Th2s we observed arose from exTregs, or Tregs that undergo reprogramming, lose Foxp3 expression, and can convert to Th2-like cells. ExTregs secrete pro-inflammatory cytokines and lose their suppressive function[52]. TCR sequence analysis, differentiation trajectory analysis, and gene expression profile all indicate that Th2 regulator cells are a unique ATT subpopulation with the capacity for immune regulatory functions. Currently, the mechanism(s) by which adipose tissue Th2 cells are differentiated and programed to express regulatory functions in lean and/or with stHFD and why Th2 cells are severely reduced with obesity remains to be elucidated. One is hypothesis is the requirement of *Pparg* to drive an adipose tissue Th2 phenotype, as be observed its expression in both Th2 and Tregs. PPAR-γ has been described as critical for adipose tissue Treg development and can promote Th2 polarization in allergy and parasite mouse models[69], however, very little is known about the requirement and function of PPAR-γ in Th2 cells in adipose tissue.

Identification of the factors that generate, activate, and sustain these cells in VAT and if these cells also reside and/or can be induced in SAT would be key in harnessing their potential therapeutic benefits in controlling adipose tissue inflammation with obesity.

We anticipated observing clonal expansion of specific ATT clones that were present in lean adipose tissue and expanded with 14d of HFD. However, our TCR sequencing analysis did not support this hypothesis, as were unable to identify a clonal match between lean and HFD fed mice. It is possible that this is a result of a limitation of the experimental design and technique, as we could not sequence the same sample of fat from the same mouse in a lean state and then after stHFD. Interestingly, we were able to observe an increase in unique TCR sequences (meaning each T cell has a TCR receptor with a sequence that different from every other T cell) with 14d of HFD in only the Th2 regulatory cells. Similarly, using the OT-II mouse model, we identified that TCRVβ5^+^ T cells proliferated the most in response to HFD, suggesting a potential role for this TCR chain in recognizing the activation signals present in the adipose tissue microenvironment. Previous reports suggest that T cell that express the TCRVβ5 chain might recognize self-antigens, as expanded usage of TCRVβ5 chains in Tregs was observed in patients with autoimmune neutropenia[70]. These data suggest activation and proliferation of these cells might be

In summary, stHFD results in the accumulation of ATTs only in male VAT, and not male iWAT or female mice. Further, ATT accumulation featured proliferation of memory, and not naïve, ATTs, suggesting a role for unique role of resident memory ATTs to respond to local adipose tissue stimuli. Of accumulating ATTs in VAT, we describe the first report of Th2s with an immunoregulatory phenotype that increase in abundance with stHFD. We also demonstrate that proliferating ATTs are associated with expression of TCRVβ5. Our study provides insights into the temporal kinetics and phenotypes of ATT with stHFD and highlight a novel underexplored ATT Th2 subpopulation with an immunoregulatory profile that serves as an attractive therapeutic target. Elucidation of the mechanism of Th2 regulatory cell induction could aid in controlling adipose tissue inflammation with obesity.

## CREDIT AUTHORSHIP STATEMENT

**Ramiah D. Jacks**: Conceptualization, Data curation, Formal analysis, Funding acquisition, Methodology, Investigation, Visualization, Writing – original draft, Writing – review & editing. **Peizi Wu**: Data duration, Formal analysis. **Jennifer L. DelProposto**: Investigation. **Carey N. Lumeng**: Conceptualization, Funding acquisition, Methodology, Visualization, Supervision, Writing – original draft, Writing – review & editing.

## DATA AVAILABLITY

UMAP and processed data can be viewed at the Single Cell Portal SCP3314.

---

**Supplemental Figure 1.**
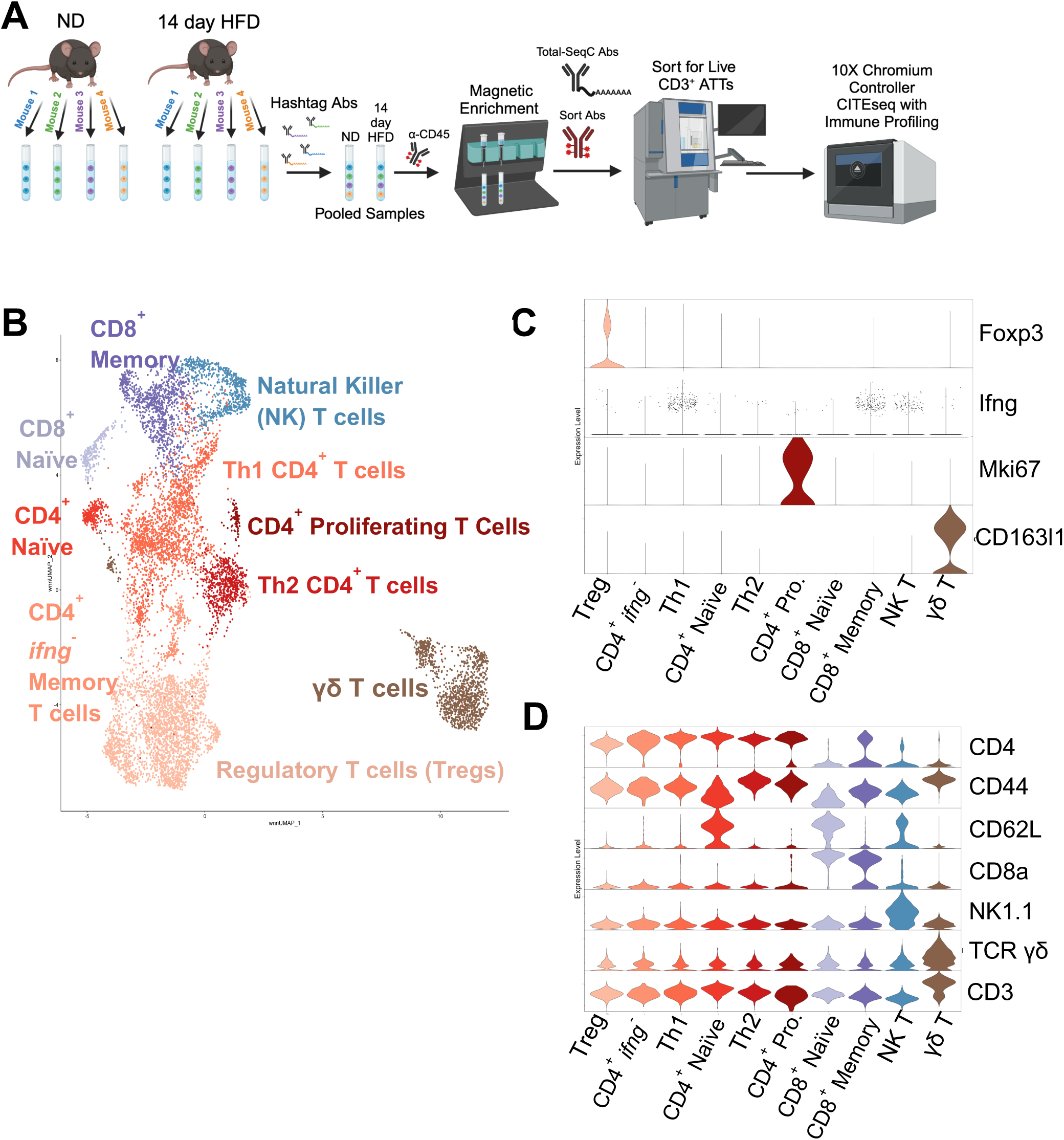
CITEseq reveals Adipose Tissue T cell populations. (A) Schematic of cellular indexing of transcriptomes and epitopes (CITE-seq) approach on CD3^+^ ATTs using hash tag antibodies (Abs) and Total-SeqC antibodies. (B) Uniform Manifold Approximation and Projection (UMAP) need to know how to describe this better. Selected markers based on (C) gene expression and (D) surface protein expression. Figure 1A was created with Biorender.

**Supplemental Figure 2.**
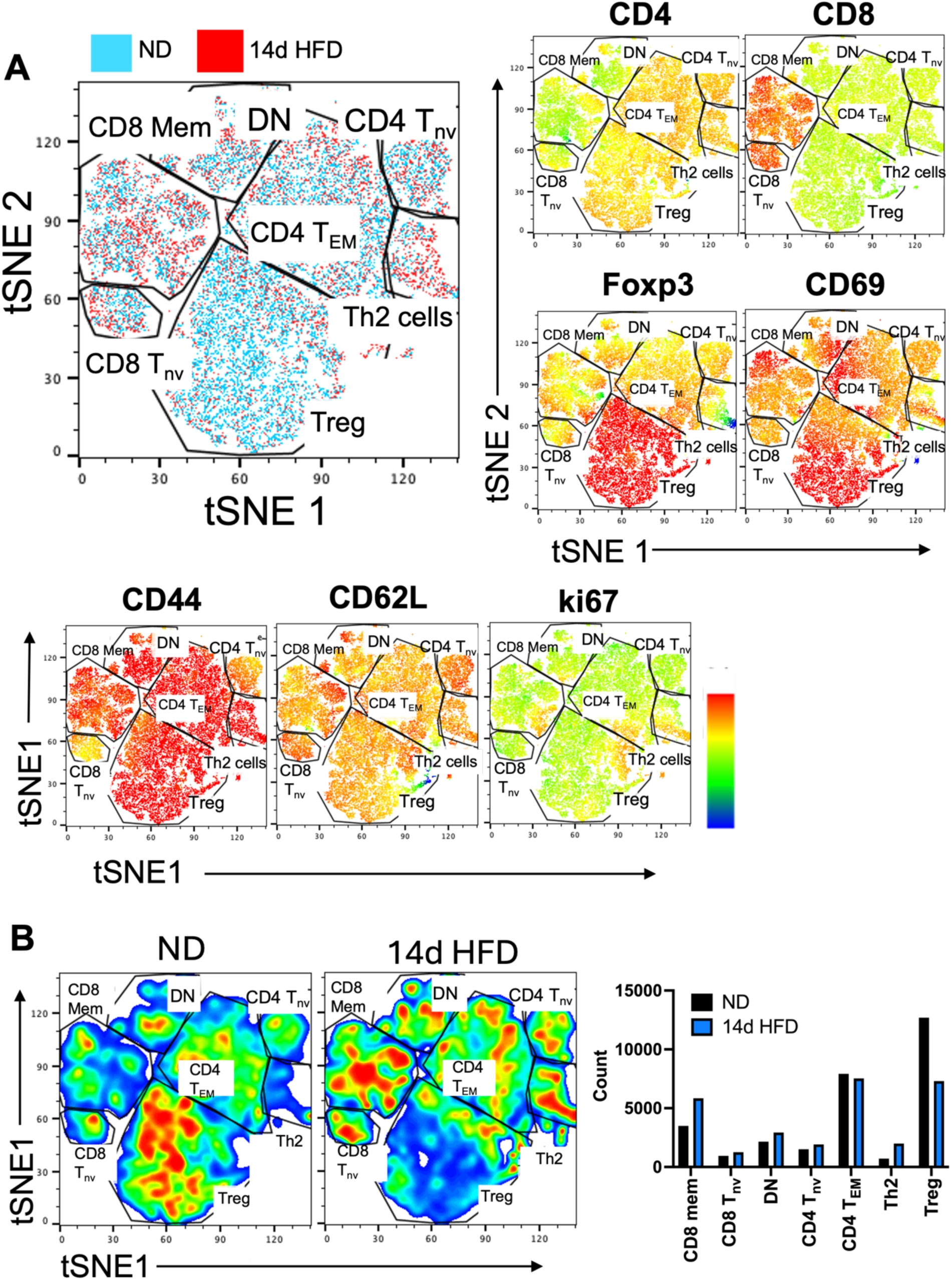
Flow Cytometry data recapitulates CITEseq Adipose Tissue T cell changes with 14 days of HFD. (A) tSNE generated from concatenate samples from ND and 14 day HFD fed mice. Clusters identified based on expression of CD4, CD8, Foxp3, CD69, CD44, CD62L and ki67. (B) Heatmap tSNE ND and 14d HFD fed mice (left) and quantitation with each cluster (right).

**Supplemental Figure 3.**
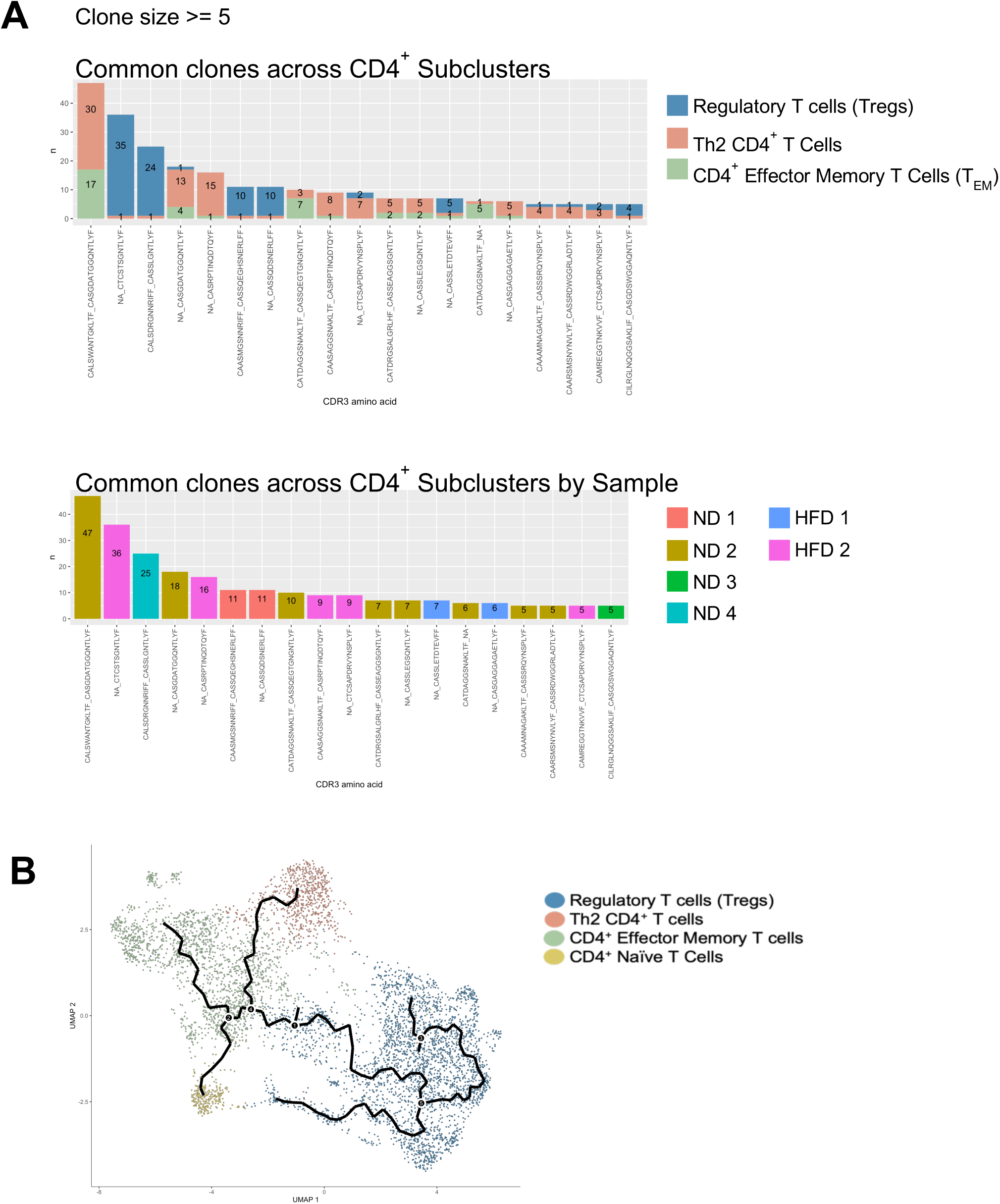
Th2 regulatory cells do not arise from ex-Tregs. (A) Number of cells with the same TCR sequence (clone) grouped by CD4+ subcluster (Treg, Th2, and T_EM_) (top) and group by individual mouse (ND 1-4 or HFD 1-2) (bottom). (B) Monocle trajectory analysis comparing the distance between Th2 regulatory cells and Tregs.

**Supplemental Table 1.** Anti-mouse antibodies used for flow cytometry analysis of Adipose Tissue T cells

**Supplemental Table 2.** Reagents used for Hashtag staining and Adipose Tissue T cell sorting

**Supplemental Table 3.** Complete List of anti-mouse antibodies in the Biolegend TotalSeq-C mouse universal cocktail used for CITESeq

**Supplemental Table 4.** RNA markers for UMAP plots of ATT cells

**Supplemental Table 5.** ADT markers for UMAP plots of ATT cells

**Supplemental Table 6.** Differentially Expressed Genes in ATT subpopulations between ND and HFD

**Supplemental Table 7.** GSEA for CD4^+^ Treg ATT Subpopulation

**Supplemental Table 8.** GSEA for γδ T cell ATT Subpopulation

**Supplemental Table 9.** GSEA for Th2 ATT Subpopulation

**Supplemental Table 10.** RNA markers for UMAP plots Gated on CD4^+^ ATTs

**Supplemental Table 11.** ADT markers for UMAP plots Gated on CD4^+^ ATTs

## REFERENCES

1. Censin, J.C., et al., Causal relationships between obesity and the leading causes of death in women and men. PLoS Genet, 2019. 15(10): p. e1008405.

2. Kawai, T., M.V. Autieri, and R. Scalia, Adipose tissue inflammation and metabolic dysfunction in obesity. American Journal of Physiology-Cell Physiology, 2021. 320(3): p. C375–C391.

3. Grant, R.W. and V.D. Dixit, Adipose tissue as an immunological organ. Obesity (Silver Spring), 2015. 23(3): p. 512–8.

4. Jacks, R.D. and C.N. Lumeng, Macrophage and T cell networks in adipose tissue. Nature Reviews Endocrinology, 2024. 20(1): p. 50–61.

5. Lu, J., et al., Adipose Tissue-Resident Immune Cells in Obesity and Type 2 Diabetes. Front Immunol, 2019. 10: p. 1173.

6. Cottam, M.A., et al., Multiomics reveals persistence of obesity-associated immune cell phenotypes in adipose tissue during weight loss and weight regain in mice. Nat Commun, 2022. 13(1): p. 2950.

7. Park, C.S. and N. Shastri, The Role of T Cells in Obesity-Associated Inflammation and Metabolic Disease. Immune Netw, 2022. 22(1): p. e13.

8. Nishimura, S., et al., CD8+ effector T cells contribute to macrophage recruitment and adipose tissue inflammation in obesity. Nature Medicine, 2009. 15(8): p. 914–920.

9. Stolarczyk, E., et al., Improved Insulin Sensitivity despite Increased Visceral Adiposity in Mice Deficient for the Immune Cell Transcription Factor T-bet. Cell Metabolism, 2013. 17(4): p. 520–533.

10. Bradley, D., et al., Interferon gamma mediates the reduction of adipose tissue regulatory T cells in human obesity. Nature Communications, 2022. 13(1): p. 5606.

11. Cho, K.W., et al., An MHC II-dependent activation loop between adipose tissue macrophages and CD4+ T cells controls obesity-induced inflammation. Cell Rep, 2014. 9(2): p. 605–17.

12. Morris, D.L., et al., Adipose Tissue Macrophages Function As Antigen-Presenting Cells and Regulate Adipose Tissue CD4+ T Cells in Mice. Diabetes, 2013. 62(8): p. 2762–2772.

13. Winer, S., et al., Normalization of obesity-associated insulin resistance through immunotherapy. Nat Med, 2009. 15(8): p. 921–9.

14. Blank, C.U., et al., Defining ‘T cell exhaustion’. Nature Reviews Immunology, 2019. 19(11): p. 665–674.

15. Porsche, C.E., et al., Obesity results in adipose tissue T cell exhaustion. JCI Insight, 2021. 6(8).

16. Painter, S.D., I.G. Ovsyannikova, and G.A. Poland, The weight of obesity on the human immune response to vaccination. Vaccine, 2015. 33(36): p. 4422–9.

17. Abd Alhadi, M., et al., Obesity Is Associated with an Impaired Baseline Repertoire of Anti-Influenza Virus Antibodies. Microbiol Spectr, 2023. 11(3): p. e0001023.

18. Jang, S., W. Hong, and Y. Moon, Obesity-compromised immunity in post-COVID-19 condition: a critical control point of chronicity. Frontiers in Immunology, 2024. 15.

19. Giamarellos-Bourboulis, E.J., et al., Complex Immune Dysregulation in COVID-19 Patients with Severe Respiratory Failure. Cell Host Microbe, 2020. 27(6): p. 992–1000.e3.

20. Muir, L.A., et al., Frontline Science: Rapid adipose tissue expansion triggers unique proliferation and lipid accumulation profiles in adipose tissue macrophages. J Leukoc Biol, 2018. 103(4): p. 615–628.

21. Hao, Y., et al., Dictionary learning for integrative, multimodal and scalable single-cell analysis. Nature biotechnology, 2024. 42(2): p. 293–304.

22. Hafemeister, C. and R. Satija, Normalization and variance stabilization of single-cell RNA-seq data using regularized negative binomial regression. Genome biology, 2019. 20(1): p. 296.

23. Aran, D., et al., Reference-based analysis of lung single-cell sequencing reveals a transitional profibrotic macrophage. Nature immunology, 2019. 20(2): p. 163– 172.

24. Heng, T.S., et al., The Immunological Genome Project: networks of gene expression in immune cells. Nature immunology, 2008. 9(10): p. 1091–1094.

25. Korotkevich, G., et al., Fast gene set enrichment analysis. biorxiv, 2016: p. 060012.

26. Yang, Q., K.R. Safina, and N. Borcherding, scRepertoire 2: Enhanced and Efficient Toolkit for Single-Cell Immune Profiling. bioRxiv, 2024: p. 2024.12. 31.630854.

27. Qiu, X., et al., Reversed graph embedding resolves complex single-cell trajectories. Nature methods, 2017. 14(10): p. 979–982.

28. Künzli, M. and D. Masopust, CD4+ T cell memory. Nature Immunology, 2023. 24(6): p. 903–914.

29. Luckheeram, R.V., et al., CD4⁺T cells: differentiation and functions. Clin Dev Immunol, 2012. 2012: p. 925135.

30. Kautzky-Willer, A., M. Leutner, and J. Harreiter, Sex differences in type 2 diabetes. Diabetologia, 2023. 66(6): p. 986–1002.

31. Goossens, G.H., J.W. Jocken, and E.E. Blaak, Sexual dimorphism in cardiometabolic health: the role of adipose tissue, muscle and liver. Nature Reviews Endocrinology, 2021. 17(1): p. 47–66.

32. Cooper, A.J., et al., Sex/Gender Differences in Obesity Prevalence, Comorbidities, and Treatment. Curr Obes Rep, 2021. 10(4): p. 458–466.

33. Varghese, M., et al., Sex Differences in Inflammatory Responses to Adipose Tissue Lipolysis in Diet-Induced Obesity. Endocrinology, 2018. 160(2): p. 293– 312.

34. Hausmann, F., et al., DISCERN: deep single-cell expression reconstruction for improved cell clustering and cell subtype and state detection. Genome Biology, 2023. 24(1): p. 212.

35. Tager, A.M., et al., Leukotriene B4 receptor BLT1 mediates early effector T cell recruitment. Nat Immunol, 2003. 4(10): p. 982–90.

36. Castro-Sánchez, P., et al., Regulation of CD4+ T Cell Signaling and Immunological Synapse by Protein Tyrosine Phosphatases: Molecular Mechanisms in Autoimmunity. Frontiers in Immunology, 2019. 10.

37. Borys, S.M., et al., The Yin and Yang of Targeting KLRG1(+) Tregs and Effector Cells. Front Immunol, 2022. 13: p. 894508.

38. McArdel, S.L., C. Terhorst, and A.H. Sharpe, Roles of CD48 in regulating immunity and tolerance. Clinical Immunology, 2016. 164: p. 10–20.

39. Joller, N., et al., Treg cells expressing the coinhibitory molecule TIGIT selectively inhibit proinflammatory Th1 and Th17 cell responses. Immunity, 2014. 40(4): p. 569–81.

40. Mehta, P., A.M. Nuotio-Antar, and C.W. Smith, γδ T cells promote inflammation and insulin resistance during high fat diet-induced obesity in mice. Journal of Leukocyte Biology, 2014. 97(1): p. 121–134.

41. Dubois, C.M., et al., Evidence that furin is an authentic transforming growth factor-beta1-converting enzyme. Am J Pathol, 2001. 158(1): p. 305–16.

42. Odagiu, L., et al., Role of the Orphan Nuclear Receptor NR4A Family in T-Cell Biology. Front Endocrinol (Lausanne), 2020. 11: p. 624122.

43. Andreotti, A.H., et al., T-cell signaling regulated by the Tec family kinase, Itk. Cold Spring Harb Perspect Biol, 2010. 2(7): p. a002287.

44. Conley, J.M., M.P. Gallagher, and L.J. Berg, T Cells and Gene Regulation: The Switching On and Turning Up of Genes after T Cell Receptor Stimulation in CD8 T Cells. Front Immunol, 2016. 7: p. 76.

45. Yukawa, M., et al., AP-1 activity induced by co-stimulation is required for chromatin opening during T cell activation. J Exp Med, 2020. 217(1).

46. Das, T., et al., A20/Tumor Necrosis Factor α-Induced Protein 3 in Immune Cells Controls Development of Autoinflammation and Autoimmunity: Lessons from Mouse Models. Front Immunol, 2018. 9: p. 104.

47. Denizot, F., et al., Novel structures CTLA-2 alpha and CTLA-2 beta expressed in mouse activated T cells and mast cells and homologous to cysteine proteinase proregions. Eur J Immunol, 1989. 19(4): p. 631–5.

48. Moreau, J.M., et al., Transforming growth factor-β1 in regulatory T cell biology. Sci Immunol, 2022. 7(69): p. eabi4613.

49. Rudensky, A.Y., Regulatory T cells and Foxp3. Immunol Rev, 2011. 241(1): p. 260–8.

50. Cipolletta, D., et al., PPAR-γ is a major driver of the accumulation and phenotype of adipose tissue Treg cells. Nature, 2012. 486(7404): p. 549–53.

51. Kanhere, A., et al., T-bet and GATA3 orchestrate Th1 and Th2 differentiation through lineage-specific targeting of distal regulatory elements. Nature Communications, 2012. 3(1): p. 1268.

52. Saxena, V., et al., Mechanisms of exTreg induction. Eur J Immunol, 2021. 51(8): p. 1956–1967.

53. Robertson, J.M., P.E. Jensen, and B.D. Evavold, DO11.10 and OT-II T cells recognize a C-terminal ovalbumin 323-339 epitope. J Immunol, 2000. 164(9): p. 4706–12.

54. Nishimura, S., et al., CD8+ effector T cells contribute to macrophage recruitment and adipose tissue inflammation in obesity. Nature medicine, 2009. 15(8): p. 914–920.

55. Liao, X., et al., Adipose stem cells control obesity-induced T cell infiltration into adipose tissue. Cell Reports, 2024. 43(3): p. 113963.

56. Han, S.J., et al., White Adipose Tissue Is a Reservoir for Memory T Cells and Promotes Protective Memory Responses to Infection. Immunity, 2017. 47(6): p. 1154–1168.e6.

57. Dhokte, S. and K. Czaja, Visceral Adipose Tissue: The Hidden Culprit for Type 2 Diabetes. Nutrients, 2024. 16(7).

58. Zhang, S., et al., The Alterations in and the Role of the Th17/Treg Balance in Metabolic Diseases. Front Immunol, 2021. 12: p. 678355.

59. Li, C., et al., Interferon-α-producing plasmacytoid dendritic cells drive the loss of adipose tissue regulatory T cells during obesity. Cell Metab, 2021. 33(8): p. 1610–1623.e5.

60. Ishikawa, A., et al., Estrogen regulates sex-specific localization of regulatory T cells in adipose tissue of obese female mice. PLoS One, 2020. 15(4): p. e0230885.

61. Varghese, M., et al., Sex hormones regulate metainflammation in diet-induced obesity in mice. J Biol Chem, 2021. 297(5): p. 101229.

62. Goldberg, E.L., et al., Ketogenesis activates metabolically protective γδ T cells in visceral adipose tissue. Nature Metabolism, 2020. 2(1): p. 50–61.

63. McLaughlin, T., et al., T-cell profile in adipose tissue is associated with insulin resistance and systemic inflammation in humans. Arterioscler Thromb Vasc Biol, 2014. 34(12): p. 2637–43.

64. Schmidt, V., et al., Obesity-Mediated Immune Modulation: One Step Forward, (Th)2 Steps Back. Front Immunol, 2022. 13: p. 932893.

65. Wang, J., X. Zhao, and Y.Y. Wan, Intricacies of TGF-β signaling in Treg and Th17 cell biology. Cellular & Molecular Immunology, 2023. 20(9): p. 1002–1022.

66. Deng, Z., et al., TGF-β signaling in health, disease and therapeutics. Signal Transduction and Targeted Therapy, 2024. 9(1): p. 61.

67. Li, Y., et al., STAT6 and Furin Are Successive Triggers for the Production of TGF-β by T Cells. The Journal of Immunology, 2018. 201(9): p. 2612–2623.

68. Lee, M.-J., Transforming growth factor beta superfamily regulation of adipose tissue biology in obesity. Biochimica et Biophysica Acta (BBA) - Molecular Basis of Disease, 2018. 1864(4, Part A): p. 1160–1171.

69. Chen, T., et al., PPAR-γ promotes type 2 immune responses in allergy and nematode infection. Sci Immunol, 2017. 2(9).

70. Goda, S., et al., Possible involvement of regulatory T cell abnormalities and variational usage of TCR repertoire in children with autoimmune neutropenia. Clin Exp Immunol, 2021. 204(1): p. 1–13.

